# Engineering a Glucose-Inducible Whole-Cell Biosensor via CRISPRi-Based Promoter Reprogramming

**DOI:** 10.64898/2026.04.09.717388

**Authors:** Mani Gupta, Anshuman Das, Satyadip Paul, Supratim Datta

**Affiliations:** Protein and Organism Engineering Laboratory, Department of Biological Sciences, Indian Institute of Science Education and Research Kolkata, Mohanpur 741246, India; Center for the Climate and Environmental Studies, Indian Institute of Science Education and Research Kolkata, Mohanpur 741246, India; Centre for Advanced Functional Materials, Indian Institute of Science Education and Research, Kolkata, Mohanpur 741246, India

## Abstract

Precise monitoring of intracellular glucose dynamics is essential for understanding carbon flux, optimizing microbial bioprocesses, and enabling responsive control of engineered metabolic pathways. Here, we develop a modular whole-cell biosensor in *Escherichia coli* that converts the native glucose repression phenotype of a CAP-sensitive promoter into a tunable, glucose-inducible output using CRISPR interference (CRISPRi). By placing a guide RNA (gRNA) under the control of the CAP promoter and positioning dCas9 to target the -10 region of a constitutive promoter driving sfGFP, we created an inversion circuit in which glucose suppresses gRNA expression, thereby relieving dCas9-mediated repression and activating fluorescence. Systematic evaluation of gRNA strand orientation and target site selection revealed that template-strand targeting yielded strong repression (∼90 %) but reduced sensing range, whereas moderately repressive non-template gRNAs (∼27-35 % repression) enabled optimal signal inversion. The resulting biosensor demonstrated a robust, linear fluorescence response across 200 μM -50 mM glucose (R² > 0.97), with high specificity against other sugars and a strong correlation between glucose consumption and fluorescence accumulation (R² ≈ 0.996). To extend the functionality of the platform, we integrated the sensor with a secreted β-glucosidase module that hydrolyzes cellobiose to glucose. The biosensor accurately quantified glucose released during cellobiose degradation, with engineered strains producing up to ∼33 mM glucose from 50 mM cellobiose in a two-plasmid system. This coupling of enzymatic conversion with intracellular sensing enabled real-time, non-destructive monitoring of metabolic transitions. Together, this work establishes a programmable CRISPRi-based strategy for inverting native promoter logic and provides a sensitive, specific, and modular platform for metabolite sensing in bacteria. The approach is broadly applicable for dynamic pathway regulation, monitoring carbon fluxes, and building responsive genetic circuits in metabolic engineering and synthetic microbial ecosystems.

TOC

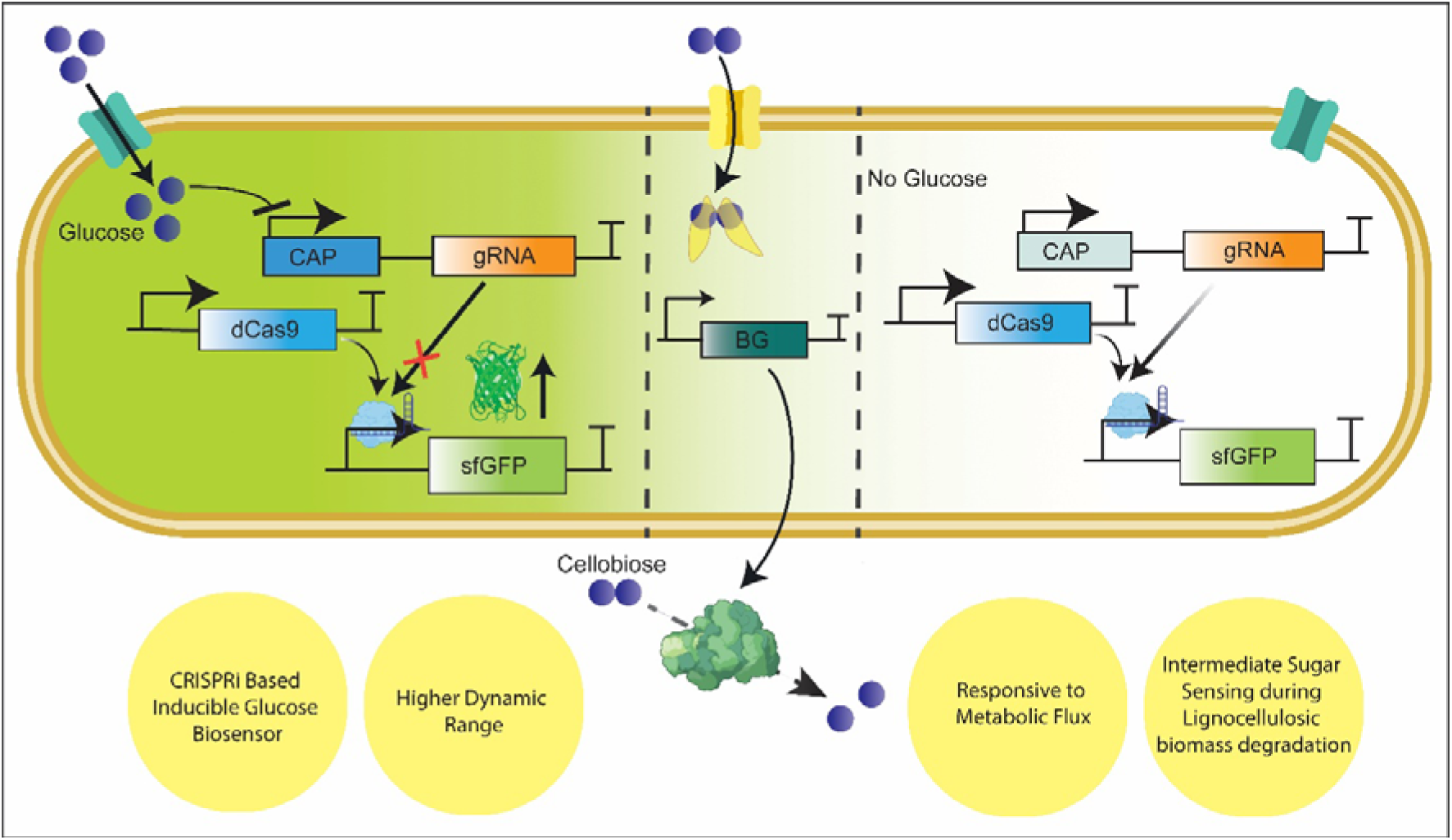

## Introduction

Whole-cell biosensors utilizing *Escherichia coli* are powerful tools for detecting and quantifying environmental and biological analytes due to their sensitivity, specificity, and real-time monitoring capabilities^1–3^. *E. coli*, a well-characterized and genetically tractable microorganism, serves as an ideal chassis for engineering synthetic biological circuits that produce easily measurable outputs, like fluorescence, luminescence, or color changes when responding to target molecules^4, 5^. These biosensors utilize natural or synthetic promoters regulated by specific inducers for sensitive detection of small molecules, toxins, heavy metals, and quorum-sensing signals^1^. Recent advancements in synthetic biology and enzyme engineering have improved the performance of *E. coli* whole-cell sensors by improving signal-to-noise ratios, expanding the detectable analytes range, and integrating multiplexed sensing capabilities^6, 7^. Innovations like modular genetic circuits, directed evolution of sensing elements, and novel reporter systems have increased robustness, dynamic range, and operational stability, broadening their applications in environmental monitoring, medical diagnostics, and industrial bioprocess control^1, 6, 8–11^. These genetically encoded systems facilitate real-time, high-throughput monitoring of metabolic states.

Conventional static regulation strategies like constitutive promoter tuning, ribosome binding site (RBS) optimization, or protein degradation tags offer the advantage of real-time responsiveness^12–14^. These sensors can dynamically adjust metabolic fluxes, limit the accumulation of toxic intermediates, and enhance yields. Various biosensors responsive to intracellular metabolites like shikimic acid^15^, lysine^16^, lactams^17^, lactic acid^18^, ethanol^19^, vanillate^20^, sugar phosphates^21, 22^ and pyruvate^23, 24^ use natural transcription factors or synthetic circuits to translate metabolite concentrations into reporter gene expression for real-time metabolic feedback regulation. However, existing biosensors often have limitations due to their reliance on specific intracellular intermediates, restricting their functionality to particular pathway nodes and hindering broader physiological sensing. This reliance can also introduce cross-regulation, complicating the modularity of synthetic systems.

Glucose, a key metabolic intermediate, is a compelling target for biosensing due to its impact on cellular energy and growth in *E. coli* and other microbes^25, 26^. In industrial biotechnology, fluctuations in glucose concentration affect fermentation performance, product yield, and cellular stress responses. Precise, dynamic, and real-time glucose sensing is crucial for optimizing engineered metabolic pathways and microbial bioprocesses. However, developing robust glucose biosensors is challenging, due to carbon catabolite repression (CCR)^25^. In *E. coli*, CCR is mediated by the catabolite activator protein (CAP) and cyclic adenosine monophosphate (cAMP)^27^. Under low-glucose conditions, there is an increase in cAMP levels, which bind to CAP and activate transcription from CAP-sensitive promoters. In contrast, high glucose levels decrease cAMP levels, reducing CAP–cAMP complex formation and suppressing transcription. Existing glucose biosensors relying on these natural repression mechanisms face limitations, such as a narrow dynamic range, lower signal output at high glucose concentrations, and complexity in high-throughput contexts^28^.

Lignocellulosic biomass, the most abundant renewable carbon resource on Earth, has over 200 billion tons of agricultural residues generated annually. Efficient saccharification of cellulose and hemicellulose into fermentable sugars, like cellobiose and glucose, is vital for sustainable biofuel and biochemical production. ^29, 30^ Real-time monitoring of intermediate and final sugar concentrations during enzymatic hydrolysis would accelerate cellulase screening, improve process control, and enhance bioconversion efficiency. However, conventional analytical methods are labor-intensive and destructive, limiting their applicability in high-throughput industrial workflows.

To tackle these challenges, synthetic biology provides novel design frameworks that integrate orthogonal control systems, such as CRISPR interference (CRISPRi). By using programmable, non-degradative repression through dCas9 and guide RNAs^31–37^, custom logic circuits with modified or tunable output behaviors can be designed. This approach can invert a CAP-regulated promoter, converting a naturally glucose-repressed system into a synthetic, glucose-inducible biosensor. Our engineered *E. coli* strain uses a dCas9–gRNA system to repress a CAP-sensitive promoter in the absence of glucose. Adding glucose reduces CAP activity and relieves dCas9 repression, triggering fluorescence that correlates with glucose concentration. The modularity, orthogonality, and programmability of our design highlight the potential of CRISPRi-driven biosensors for real-time metabolite detection and dynamic pathway regulation. This study demonstrates the application of this approach for glucose sensing in *E. coli*, overcoming the limitations of traditional repression-based systems and enabling sensing of intermediate sugars such as cellobiose during saccharification. To extend this framework for lignocellulosic process monitoring, a cellobiose-responsive module based on CelR, a GalR–LacI family transcriptional regulator that binds a palindromic operator and is derepressed upon cellobiose binding, was integrated. This hierarchical design enables enzymatic conversion of cellobiose to glucose and simultaneous quantification of the resulting metabolic intermediate within a single engineered strain.

## Materials and Methods

### Strains, Plasmids, Primers, and Growth Medium

*Escherichia coli* DH5α was used for plasmid construction and cloning, while *E. coli* BL21(DE3) was used for protein expression and functional assays. All strains, plasmids, and primers used in this study are listed in Tables S1 and S2, and the annotated sequences of synthetic genetic elements are provided in Table S3. Plasmid transformations were performed using a standard heat-shock protocol with chemically competent cells prepared in-house. Bacterial cultures were grown in Luria-Bertani (LB) broth or M9 minimal medium, supplemented with appropriate antibiotics according to the selection marker of each plasmid.

### Plasmid construction

All plasmids were assembled using a combination of EcoFlex^38^ Modular Cloning (MoClo) and Gibson Assembly. In the EcoFlex system, standardized genetic parts (promoters, ribosome-binding sites (RBSs), open reading frames (ORFs), and terminators) were first stored in Level□0 vectors. Individual transcription units (TUs) were assembled into Level 1 vectors using one-pot Golden Gate reactions. Multi-gene constructs were generated by assembling Level□1 TUs into Level□2 vectors via Gibson Assembly.

### Construction of CAP sensitive promoter plasmid for characterization

The CAP-binding site and core promoter were amplified from the iGEM part BBa_J04450 and cloned into the pBP-lacZ Level 0 backbone. A Level 1 transcription unit consisting of the CAP promoter, pETRBS, sfGFP as the reporter, and BBa_B0015 terminator was assembled and inserted into the pTU1-A vector (pMB1 origin, ampicillin resistance) using EcoFlex Golden Gate Assembly.

### Construction of gRNA library for CRISPRi repression and characterization

To evaluate CRISPRi-mediated repression, a guide RNA (gRNA) targeting the -10 non-template region of the J23108 promoter was cloned into pBP-ORF using primers 17 and 18 (Table S2). The corresponding level 1 gRNA transcription unit was assembled and integrated into a Level 2 construct containing dCas9 and the sfGFP reporter via Gibson assembly. Additional gRNAs targeting alternative strands and positions within the promoter or coding region were directly introduced into Level 2 constructs using Gibson Assembly with the backbone primer pair L4440 FP and L4440, along with gRNA-specific Gibson-compatible primers listed in Table S2. This two-part assembly approach allowed rapid construction and screening of multiple repression constructs. gRNAs were checked for homology with the *E. coli* genome and the plasmid sequence for off-target sites.

### Construction of the Glucose-Sensing Unit

Functional gRNA constructs identified from repression screening were incorporated into the glucose-inducible biosensor by replacing the IPTG-inducible T7 promoter upstream of the gRNA cassette with the CAP-sensitive promoter. This substitution was performed via Gibson Assembly using insert-specific primers 22-23 and vector backbone primers 24-25 (Table S2).

### Construction of the β-glucosidase Secretion and Cellobiose-Sensing Unit

Individual Level 1 transcription units, such as J23100-pETRBS-CelR-BBa_B0015, mTRC-Tag linker RBS-AnsB-BG-BBa_B0015, and mTRC-pETRBS-mCherry_ssrA-BBa_B0015 were each cloned into pTU1-A backbone. These parts were PCR-amplified with primers 26-31 (Table S2), with primer 31 serving as a common reverse primer for both 2-TU and 3-TU Level 2 assemblies. The recipient vector pTU2-a was amplified using primers 32 and 33. PCR products were purified using the QIAGEN Gel Extraction Kit and assembled into Level 2 constructs via Gibson Assembly.

A multi-unit construct containing Level 1 transcription units for CelR, mTRC-driven AnsB-BG, CAP-promoter-driven gRNA (−10/108 NT), J23100-dCas9, and J23108-sfGFP was generated by PCR amplification with primers 26-31 and 42-43, followed by Gibson Assembly into pTU2-a (pMB1 origin, chloramphenicol resistance). To enable co-transformation and ensure plasmid compatibility, the CelR-BG-CAP-gRNA cassette was cloned into pTU3-A (containing a high-copy pMB1 origin, and ampicillin resistance), while the dCas9-sfGFP cassette was assembled into pTU2-a, containing a low-copy p15A origin and chloramphenicol resistance.

### Optical density and fluorescence endpoint and Kinetic measurements

Single colonies were grown overnight in 5 mL LB medium containing the appropriate antibiotic (25 mg·L□¹ chloramphenicol or 100 mg·L□¹ ampicillin) at 37 °C with shaking at 200 rpm. Overnight cultures were diluted 1:100 into 1 mL of M9 minimal medium supplemented with the same antibiotic and the desired differential carbon source (glucose or cellobiose), and incubated in 24-well plates under identical conditions. For CRISPRi repression assays, IPTG (1-2 mM) was added after about 2 hours of growth; for β-glucosidase induction, cellobiose (0-1% w/v) was added at the same time point. Cell growth was monitored by OD□□□, sfGFP fluorescence was measured (excitation 485 nm, emission 510 nm) and mCherry fluorescence (excitation 585 nm, emission at 610 nm) was measured using a Cytation 5 multimode microplate reader (BioTek Instruments). Endpoint measurements were taken at 8 hours post-inoculation, while kinetic measurements were collected over 10-24 hours. Data were normalized to OD_600,_ where stated.

### Biochemical Glucose Quantification Assays

Glucose concentrations were quantified with a Glucose Oxidase-Peroxidase (GOD-POD) assay kit (Sigma-Aldrich). After overnight growth of *E. coli* strains carrying the glucose-sensing plasmid in LB with chloramphenicol, cultures were diluted 1:100 into M9 medium containing varying glucose levels. 1 mL cultures were taken at 2, 4.5, 6, and 8□hours post inoculation, centrifuged (13□000□rpm, 10□min, 4□°C), and the supernatants were mixed with GOD-POD reagent (40□µL diluted supernatant□+□80□µL reagent), incubated at 37□°C for 30□mins, then stopped with 80□µL 12□N H_2_SO_4_. Absorbance at 527□nm was measured on a SpectraMax 96-well reader (25□°C). Each condition was assayed in triplicate, and a standard curve generated from known glucose concentrations in M9 medium was used to calculate sample glucose levels. These concentrations were correlated with sfGFP fluorescence (normalized to OD_600_) measured at the same time points to determine glucose consumption rates and the corresponding rate of fluorescence induction.

### Quantification of β-Glucosidase Secretion via Activity Assay

To assess the secretion and enzymatic activity of β-glucosidase (BG), E. coli colonies harboring the construct J23100-CelR-mTRC-AnsB-BG were grown overnight in 5 mL LB medium containing 25 mg·L□¹ chloramphenicol at 37□°C with shaking at 200 rpm. Secondary cultures (1 mL) were inoculated at a 1:100 dilution into 24-well plates containing fresh LB medium with the same antibiotic and grown under identical conditions. After 2 hours, when the cultures reached the mid-log phase (as monitored by OD600), cellobiose was added at final concentrations of 0.1% and 1% (w/v) to induce BG expression. Culture samples were collected at 2, 4, 6, 8, 10, 12, and 14 hours post-induction. Samples were centrifuged at 13,000 rpm for 10 minutes at 4°C, and the resulting supernatants were assayed using *p*-nitrophenyl-β-D-glucopyranoside (*p*NPGlc) as substrate. For each assay, 50 µL of supernatant was mixed with 20 mM *p*NPGlc and McIlvaine buffer (pH 6.0) to a final volume of 100 µL, and incubated either at 74°C for 5 minutes (enzyme-optimal temperature) or at 37 °C for 3 hours (physiological condition).

Reactions were stopped with 100 µL 0.4 M glycine (pH 10.8), and the absorbance at 405 nm was recorded on a SpectraMax plate reader. Samples were diluted 20-fold prior to measurement to remain within the linear range. All assays were performed in triplicate.

### Immunoblotting and secretion analysis of β-glucosidase

*E. coli* BL21(DE3) cultures harboring the expression construct were grown in LB medium in shake flasks at 37□°C. When the OD600 reached approximately 0.6, expression was induced with 0.1% or 1% (w/v) cellobiose. Supernatant was collected at 2, 4, 6, and 8 hours post-induction for both enzymatic activity assays and secretion analysis. For immunoblotting, culture samples were centrifuged at 13,000 rpm for 10 minutes at 4□°C, and the resulting cell-free supernatants were run on SDS-PAGE, transferred to PVDF membranes, and probed with an anti-His monoclonal antibody followed by chemiluminescent detection.

### Cellobiose Sensing Kinetics and Cellobiose to Glucose Conversion

Individual *E. coli* colonies were initially cultured overnight in LB medium supplemented with the appropriate antibiotics (ampicillin or chloramphenicol) at 37□°C with shaking at 200 rpm. Secondary cultures were inoculated into M9 minimal medium containing the same antibiotics and varying concentrations of cellobiose, using a 1:100 dilution ratio. Cultures were grown in 24-well plates, and optical density (OD□□□) and sfGFP fluorescence (excitation: 485 nm; emission: 510 nm) were measured on a Cytation 5 multimode plate reader (BioTek) over a 20-hour period. A control strain expressing sfGFP under the J23108 promoter without any β-glucosidase or sensing elements was run in parallel to establish baseline fluorescence. To estimate the efficiency of cellobiose-to-glucose conversion, the same assay procedure was repeated with equivalent glucose concentrations instead of cellobiose (see Figures S4 and S5). The resulting fluorescence data from the glucose control experiments were used to generate a calibration curve.

## Results and Discussion

### Characterization of a CAP-Sensitive Promoter in Response to Glucose Levels

To study how a native CAP-dependent promoter responds to glucose, we constructed a reporter plasmid in *E. coli*. The plasmid contains a CAP binding site upstream of the core promoter, a strong synthetic ribosome binding site (pETRBS), sfGFP reporter gene (Figure 1, inset) and the standard transcriptional terminator, BBa_B0015. In *E. coli*, the CAP system is controlled by intracellular cyclic AMP (cAMP) levels, which are inversely correlated with glucose availability. As illustrated in Figure 1A, under low-glucose conditions, elevated cAMP levels promote the formation of a CAP–cAMP complex that binds to the CAP site upstream of the promoter. This interaction facilitates the recruitment of RNA polymerase and enhances transcription of the downstream gene (Figure 1A). Conversely, high-glucose conditions inhibit adenylate cyclase activity, leading to reduced cAMP production and diminished CAP-cAMP complex formation (Figure 1B). This inhibition results in decreased transcriptional activity, yielding only basal-level expression of the reporter gene.

**Figure 1.**
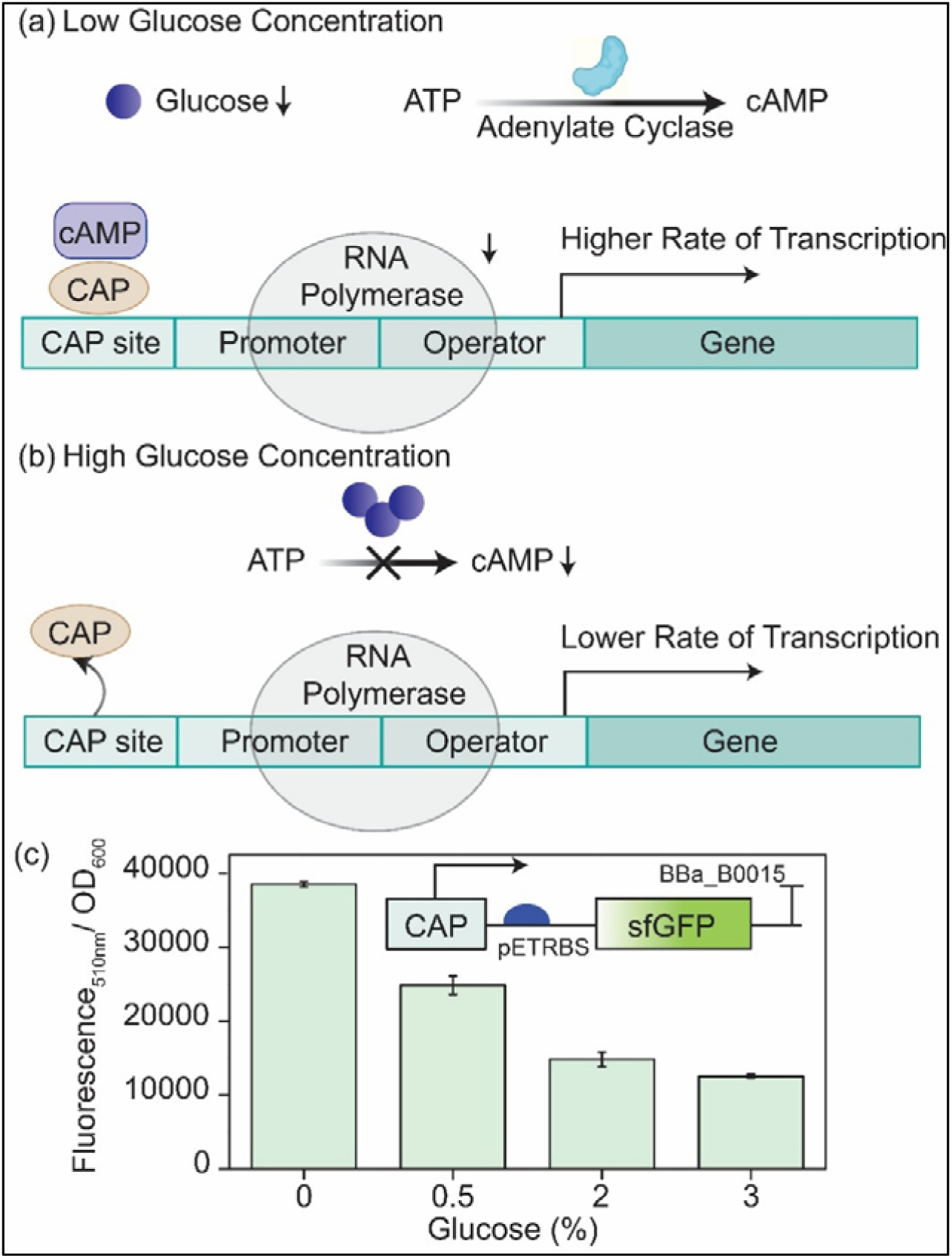
Characterization of a CAP-sensitive promoter in response to glucose levels in E. coli. (A) Schematic illustration of the native CAP–cAMP regulatory mechanism under low and high glucose conditions. Under low glucose, intracellular cAMP levels increase due to active adenylate cyclase, forming a CAP–cAMP complex that binds upstream of the promoter to enhance RNA polymerase recruitment and drive gene transcription. In contrast, high glucose inhibits cAMP synthesis, reducing CAP–cAMP complex formation and resulting in transcriptional repression. (B) Experimental validation of the CAP promoter activity using a synthetic reporter construct (inset: genetic circuit with CAP sensitive promoter, pET RBS, sfGFP, and BBa_B0015 terminator). Fluorescence output (normalized to OD_600_) was measured in cells grown with 0, 0.5, 2, and 3% glucose. A clear inverse correlation between glucose concentration and GFP expression was observed, confirming glucose-dependent repression of the CAP-sensitive promoter. Error bars represent the standard deviation from biological triplicates.

To validate this regulatory mechanism, we cultured the CAP promoter-sfGFP reporter strain in LB media supplemented with 0% (no glucose), 0.5% (moderate glucose), 2%, and 3% (high glucose). Fluorescence measurements normalized to optical density (OD) indicated that cells without glucose showed the highest fluorescence (Figure 1C), consistent with maximal CAP-cAMP complex formation and promoter activation. At 0.5% glucose, fluorescence intensity decreased significantly, indicating partial promoter repression. At 2% and 3% glucose, fluorescence levels remained low, indicating strong catabolite repression and minimal CAP binding. These observations confirm that the promoter is strongly repressed by glucose in a concentration-dependent manner, consistent with canonical CAP-mediated transcriptional control. This establishes a clear inverse correlation between extracellular glucose concentration and reporter gene expression. However, the inherent repression limits the potential of biosensor applications that require a positive correlation between the signal and a wider range of glucose levels for real-time tracking or control feedback.

### Design and Evaluation of CRISPRi-Mediated Repression at Multiple Target Sites

To synthetically invert the transcriptional output of the CAP-sensitive promoter in response to glucose, we evaluated the efficiency of CRISPR interference (CRISPRi) by directing dCas9 to various regulatory and coding regions within a modular reporter construct. Guide RNAs (gRNAs) were designed to target two primary regions: (1) the -10 region upstream of the transcription start site in two distinct promoters, J23108 and SJM910, and (2) selected regions within the coding sequence of the sfGFP gene (Figure 2A, 2C) with no off-targets. The gRNAs were placed under the control of an IPTG-inducible T7 promoter, while dCas9 was expressed constitutively, and sfGFP expression was driven by the respective target promoter (Figure 2B).

**Figure 2.**
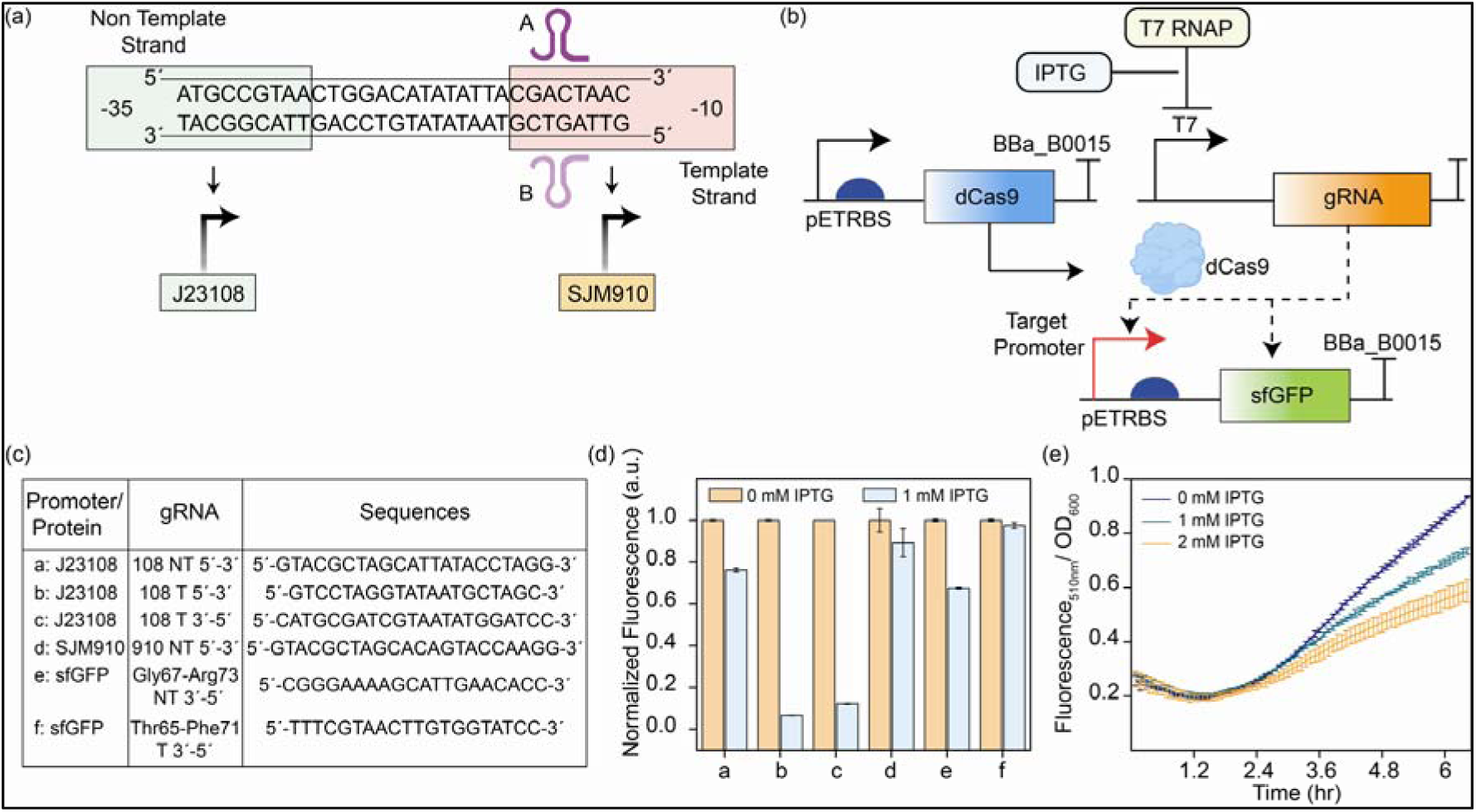
Evaluation of CRISPRi-mediated repression at multiple target sites within the reporter construct. (A) Schematic representation of the gRNA targeting two different promoters: J23108 and SJM910 (B) Schematic representation of the functioning of dCas9 and gRNA system targeting the promoter and coding region of the reporter gene. (C) Table containing different gRNA sequences targeting different sites of promoter and reporter gene. (D) Quantitative analysis of fluorescence output (normalized RFU/OD600) from *E. coli* cells transformed with each targeting construct. All gRNA-targeting strains show significant repression relative to the non-targeting control, with the promoter-targeted gRNA achieving the strongest repression. Error bars represent standard deviation from biological triplicates. (E) Kinetics of the chosen gRNA targeting -10 non-template region of J23108 promoter under IPTG inducible T7 promoter up to 8hours post induction. Expression of dCas9 is constitutive. This shows gradual reduction in Fluorescence/OD with increasing IPTG induction (1 mM, 2 mM).

Among the designs, the gRNA designated gRNA -10/108 NT, which targets the non-template strand of the -10 region in the PJ23108, demonstrated the modest repression (∼27%), while gRNA targeting the same region of SJM910 shows very low repression (∼12%). The gRNA designated gRNA -10/108T, which targets the template strand of the -10 region in the PJ23108, exhibited the strongest repression (∼94%). In all these constructs, the dCas9 handle was oriented toward the 3′ end in an inverted configuration. We also targeted template strand of the –10 region in the PJ23108 with the dCas9 handle oriented towards the 5′ end in significant repression (∼88%).

To explore the effects of CRISPRi beyond transcription initiation, we designed gRNAs targeting the non-template strand of the sfGFP coding region at two loci: Thr65-Phe71 and Gly67-Arg73. The sfGFP fluorescence was due to the beta barrel structure, mainly due to Thr65, Ser66, Gly67, hence, we hypothesized that targeting this region might reduce fluorescence by interfering with transcriptional elongation or mRNA stability The gRNA targeting Gly67-Arg73 showed moderate repression (∼33%), while the one targeting Thr65-Phe71 resulted in minimal repression (∼4%) (Figure 2D). In these constructs, the dCas9 handle was oriented toward the 3 ′ end in an inverted configuration. Fluorescence measurements revealed that all gRNAs significantly repressed sfGFP expression compared to uninduced gRNA expression control, but the degree of repression varied depending on the target site (Figure 2D). These results suggest that CRISPRi can interfere not only with transcription initiation but also with transcriptional elongation or mRNA processing. However, the degree of repression from coding region targeting was modest compared to promoter-directed gRNAs, indicating that initiation blocking strategies remain more effective for strong repression.

Variability in repression levels can be attributed to differences in gRNA binding strength at the target site. Promoter-targeting gRNAs produced the strongest repression, likely due to direct interference with RNA polymerase binding or promoter escape. Comparing all these constructs, the one that exhibited only ∼30% repression performed better, offering valuable insights into the balance between repression strength and signal inversion for dynamic sensing performance. Interestingly, while gRNA -10/108T yielded the highest repression efficiency (∼90%), its dynamic range as a glucose sensor was limited, likely due to strong baseline suppression (Supplementary Figure S6). This trade-off suggests that while tight repression is desirable for gene silencing, it may not be optimal for sensing applications that require a tunable signal response across a metabolite gradient. To observe the trend in a time responsive manner, fluorescence kinetic studies of the gRNA library strongly support the repression analysis (Figure 2E, Figure S1). These outcomes highlight the versatility of CRISPRi for programmable transcriptional repression in *E. coli* and provide critical insights into optimal gRNA positioning for achieving maximal regulatory effects. Based on our findings, we selected the gRNA targeting the PJ23108 template strand for integration into the subsequent glucose-responsive circuit.

### CRISPRi-Mediated Inversion of CAP Promoter Enables Glucose-Inducible Fluorescent Output

We constructed a synthetic gene circuit to demonstrate inversion of the CAP-sensitive promoter. The circuit contained a gRNA targeting the -10 region of the J23108 promoter, expressed under a CAP-sensitive promoter, dCas9 under a constitutive promoter, and sfGFP expression under PJ23108 (Figure 3A). Under low-glucose conditions, CAP binds upstream of the gRNA promoter, driving strong gRNA expression and recruiting dCas9 to the -10 region of the PJ23108, thereby repressing sfGFP expression. Conversely, under high-glucose conditions, CAP doesn’t form a complex due to reduced cAMP levels, leading to decreased gRNA production and restoring sfGFP expression by relieving dCas9-mediated repression. Our design effectively converts a native glucose-repressed system into a glucose inducible one.

**Figure 3.**
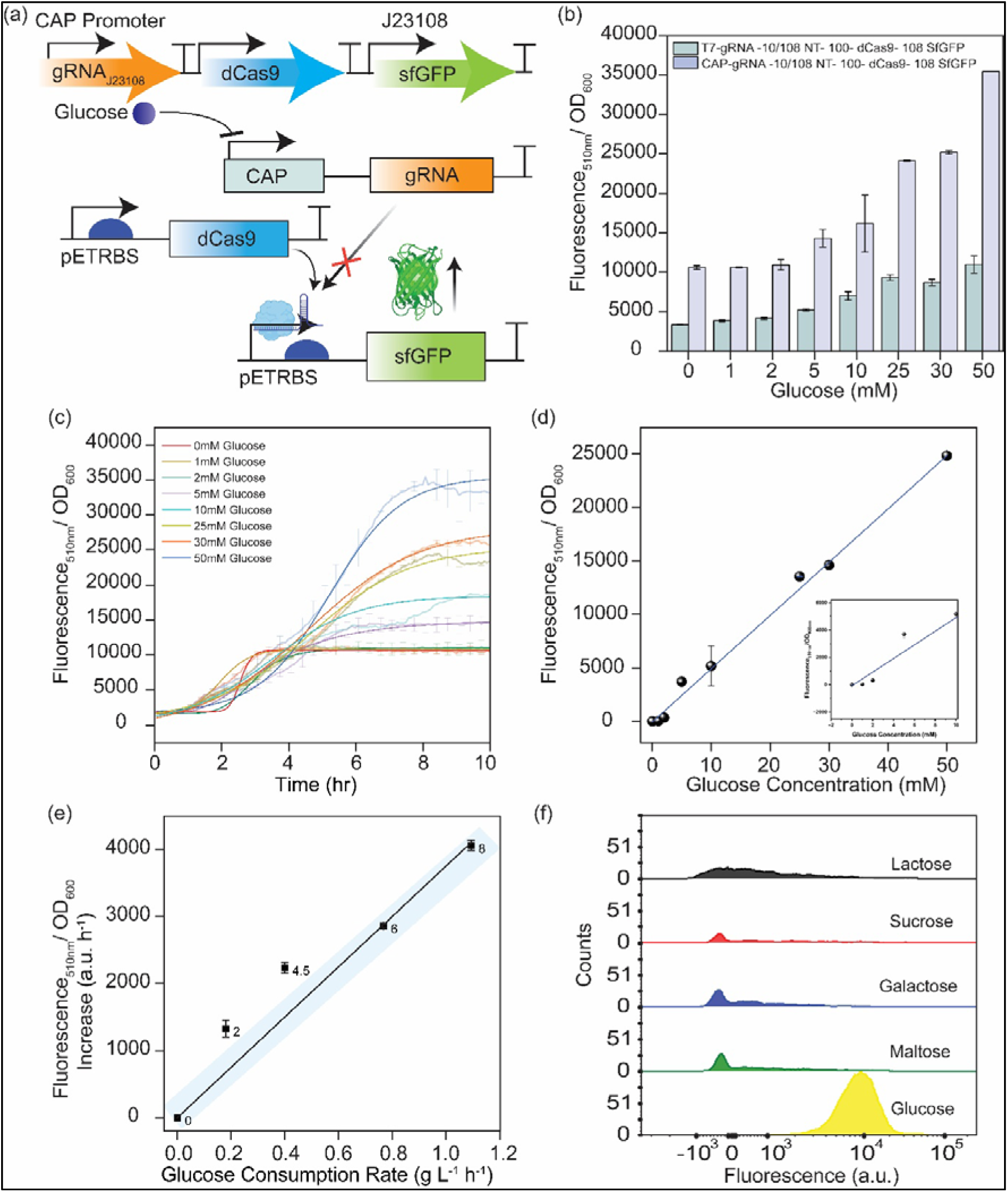
Glucose-inducible gene expression via CRISPRi-mediated inversion of CAP promoter logic. (A) Schematic of the synthetic circuit: gRNA targeting the –10 region of the J23108 promoter is expressed from a CAP-sensitive promoter, dCas9 is constitutively expressed, and sfGFP is under J23108 control. Under low glucose, active CAP promotes gRNA transcription, enabling dCas9-mediated repression of sfGFP. Under high glucose, gRNA expression is reduced, relieving repression and allowing sfGFP expression. (B) Fluorescence response (RFU/OD600) across increasing glucose concentrations (0–50 mM). A strong positive correlation was observed, indicating glucose-inducible output. Basal level of fluorescence response of a control circuit with gRNA under T7 promoter, dCas9 under constitutive promoter and sfGFP under PJ23108 promoter under similar conditions. (C) Time-course fluorescence measurement over 10 hours at varying glucose concentrations. Reporter expression increases with glucose in a dose-dependent manner. Error bars represent standard deviation from biological triplicates. (D) A linearly increasing Fluorescence (RFU/OD600) response of the Glucose Sensing circuit with increasing Glucose concentration. Error bars represent standard deviation from biological triplicates. (E) Relationship between the Fluorescence (RFU/OD600) Increase rate (per hr) and glucose consumption rate (g/L/hr) at different time points of the cell culture for the Glucose Sensing Circuit. The error bars represent standard deviation from biological triplicates. (F) Fluorescence Response of the sensing circuit to different carbon sources in flow cytometry. This shows the specificity of our circuit to glucose over other sugars.

Quantification of fluorescence across a glucose concentration range (0 to 50 mM) revealed a dose-dependent increase in sfGFP expression (Figure 3B). Basal fluorescence was recorded at 0 mM glucose, consistent with maximal gRNA-mediated repression. A sharp increase in fluorescence was observed between 1 and 10 mM glucose, demonstrating the system’s sensitivity to physiologically relevant glucose concentrations. The fluorescence signal saturated at 50 mM, demonstrating robust dynamic range and strong correlation between glucose input and reporter output.

Kinetic analysis over 10 hours further confirmed this dynamic behavior (Figure 3C). In the absence of glucose, fluorescence remained low, showing sustained CRISPRi-mediated repression. At intermediate glucose levels (2-10 mM), fluorescence gradually increased, with differences in induction onset and slope reflecting glucose-dependent gRNA repression relief. Higher concentrations (25-50 mM) led to rapid and maximal reporter expression, consistent with minimal gRNA production under suppressed CAP activity. A linear relationship was observed between the final fluorescence/OD_600_ and glucose concentration (R² = 0.99) up to 50 mM (Figure 3D), establishing the system’s quantitative dynamic range. Notably, signal saturation beyond this point was not observed under these experimental conditions. Dose-response curve analysis revealed that the biosensor achieves a limit of detection as low as 200 µM (Figure S8).

To assess the biosensor’s responsiveness in dynamic metabolic conditions, time-resolved measurements of glucose consumption were conducted alongside fluorescence accumulation (Figure 3E). Culture supernatants were collected at 2, 4.5, 6, and 8 hours for biochemical glucose quantification using a GOD-POD assay. The calculated glucose consumption rate (g L^-1^h^-1^) correlated with the rate of fluorescence/OD□□□ increase (R² = 0.996), supporting the system’s ability to report metabolic fluxes in real time. Furthermore, specificity assays confirmed that the circuit selectively responds to glucose and does not show significant activation with other common carbon sources such as lactose, sucrose, maltose, or galactose (Figure 3F), highlighting its suitability for metabolic monitoring in complex carbon backgrounds.

These results confirm our synthetic biosensor as a reliable, tunable, and metabolically responsive glucose reporter. It distinguishes between transient and sustained glucose uptake, as evidenced by the plateau in glucose consumption beyond 50 mM, despite the continued increase in sfGFP signal, likely due to intracellular retention or delayed feedback effects in metabolic processing (Table S4, Figure S9). These data confirm successful inversion of the CAP promoter through CRISPRi-mediated interference. By reprogramming CAP-mediated repression through CRISPR interference, our system generates a glucose-inducible output with an extended dynamic range and improved signal resolution compared to native repression-based designs. The modular strategy enables quantitative glucose tracking and can be readily integrated into metabolic feedback circuits or bioprocess monitoring platforms.

### Application of CRISPRi-Based Glucose Biosensor for Monitoring Cellobiose Conversion by Secreted **β**-glucosidase

We tested the glucose-biosensor by coupling it to a secreted β-glucosidase (BG) engineered to hydrolyze cellobiose to glucose. The N-terminus of the BG was fused to a secretion tag and expressed under the control of a cellobiose-inducible mtrC promoter regulated by the CelR transcription factor (Figure 4A). To report promoter activity, a second construct expressed mCherry under the same mtrC promoter. The mCherry fluorescence increased proportionally with cellobiose (0.05-1% w/v), indicating induction of PmtrC and likely co-expression of the secreted BG (Figure 4B; Figure S4). However, chromogenic BG assays in culture supernatants using *p*-nitrophenyl-β-D-glucopyranoside (*p*NPGlc) showed higher activity with 0.1% cellobiose induction than with 1 %, both under optimal BG assay conditions (74 °C for 5 minutes) and standard culture conditions (37°C for 3 hours) (Figure 4D, E).

**Figure 4.**
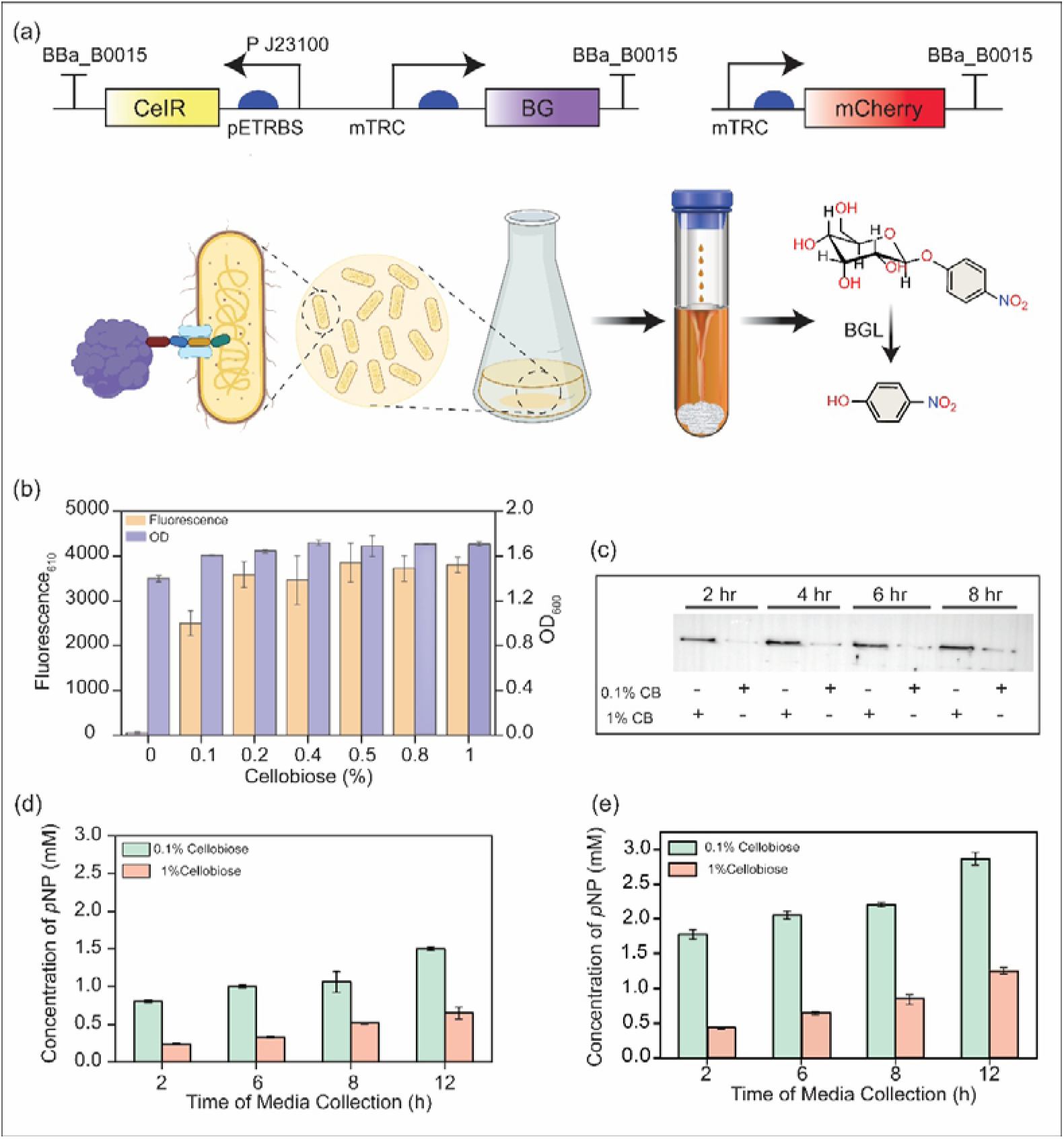
Application of CRISPRi-based glucose biosensor to detect glucose generated by secreted β-glucosidase. (A) Schematic of experimental design: Secretory-tagged β-glucosidase (BG) is expressed under the cellobiose-inducible promoter (PmtrC) regulated by CelR. mCherry is co-expressed under the same promoter as a reporter for promoter activity. Secreted BG hydrolyzes cellobiose to glucose. (B) Fluorescence and OD600 of mCherry under increasing cellobiose concentrations confirm promoter activity. (C) Western blot showing higher levels of secreted BG in 1% cellobiose induction across time points (2, 4, 6, 8 h). Enzyme activity assay at optimum temperature 74^°^ for 5min (D) and 37° and 3hours (E) using *p*NPGlc reveals higher activity at 0.1% than at 1% cellobiose, likely due to substrate competition or prior consumption of cellobiose.

To investigate the differences between reporter expression and enzymatic activity, we analyzed culture supernatants from cells induced with with 0.1% and 1% cellobiose, collected at 2, 4, 6, and 8 hours post-induction. Western blots showed greater amounts of secreted BG in the 1 % cellobiose condition at all time points, confirming that the reduced *p*NPGlc activity at 1% cellobiose induction was not due to lower BG secretion (Figure 4C). We attribute the apparent lower activity to substrate competition and consumption in the culture medium. High concentrations of cellobiose in the 1 % condition likely compete with *p*NPGlc for BG binding and/or are partially hydrolyzed to glucose prior to the assay, turnover at the BG active site during the assay, and substantial cellobiose may have both of which could reduce measurable pNPGlc turnover in endpoint assays.

### Integrated cellobiose-to-glucose conversion and sensing

We constructed a single genetic circuit that couples enzymatic cellobiose hydrolysis to the intracellular sensing of a CRISPRi-based glucose biosensor (Figure 5A). This circuit included a CelR-regulated mtrC-driven secretion module expressing AnsB-tagged β-glucosidase (BG), alongside a CAP-sensitive promoter driving gRNA expression targeting the PJ23108–sfGFP reporter, with dCas9 expressed constitutively. The use of CelR, a cellobiose-responsive transcriptional repressor, enables selective induction (Figure S3) while remaining orthogonal to glucose-mediated catabolite repression^39^ and the use of AnsB tag allows secretion of β-glucosidase (BG) into the extracellular medium^40^.

**Figure 5.**
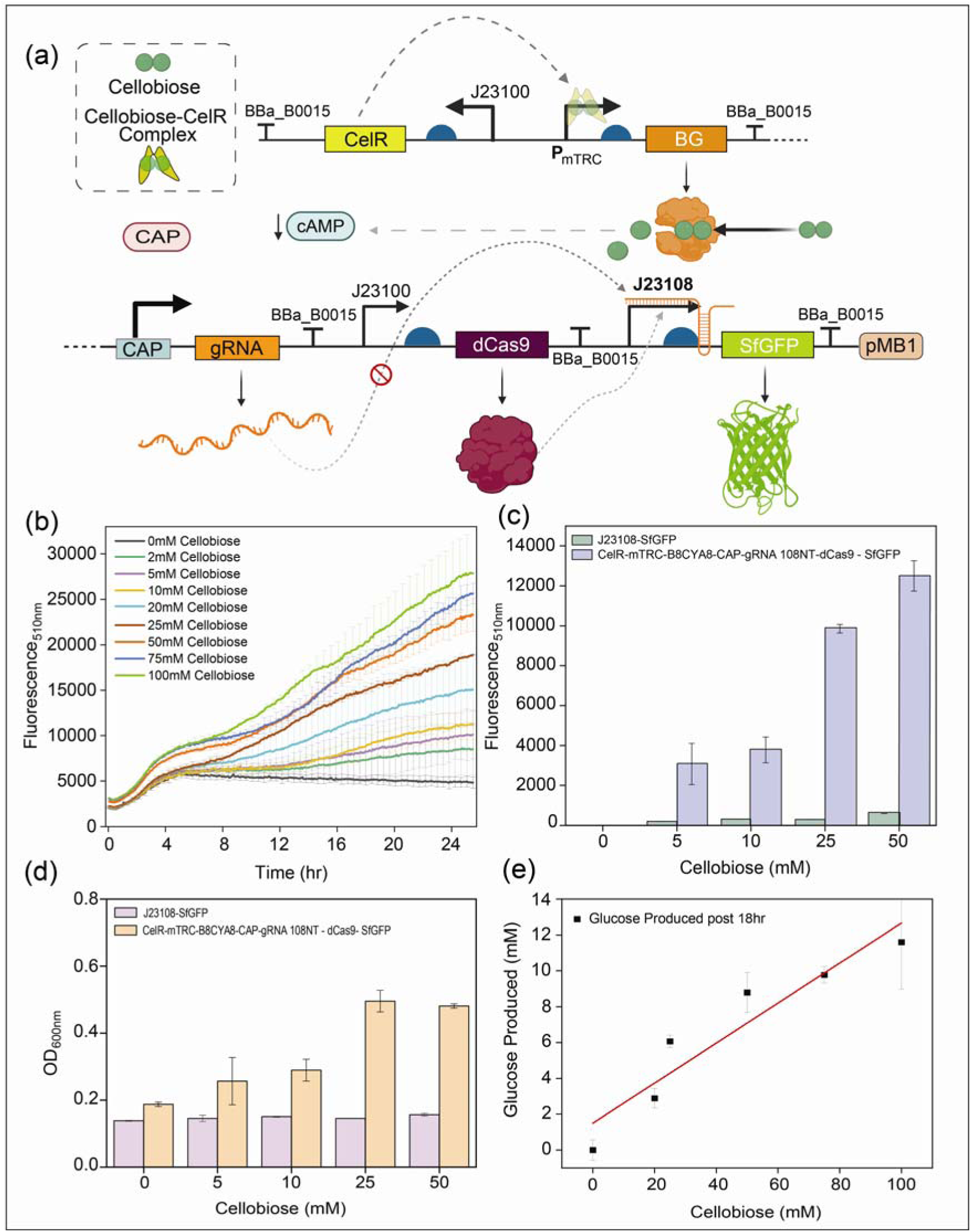
Coupling of secretory β-glucosidase and CRISPRi-based biosensor for autonomous glucose production and detection from cellobiose. (A) Schematic of the integrated circuit: CelR induces expression of secretory BG in response to cellobiose; liberated glucose represses gRNA production (via CAP promoter), relieving CRISPRi repression of sfGFP and enabling fluorescence output. (B) Time-course fluorescence confirms cellobiose-dependent induction of sfGFP signal, consistent with CRISPRi circuit logic. (C) Fluorescence measurements after 24 h in M9 + cellobiose: only strains expressing BG show signal. (D) OD600 measurements showing growth only in the BG-expressing strain, confirming glucose-supported growth. (E) Calibration curve used to estimate glucose concentration based on fluorescence output (R² = 0.91), indicating effective cellobiose-to-glucose conversion.

A control strain lacking the BG cassette but retaining the sfGFP sensor showed no growth or fluorescence under all tested conditions (Figures 5B, 5C), confirming that *E. coli* cannot natively metabolize cellobiose. Engineered strains, by contrast, displayed dose-dependent increases in both OD_600_ and sfGFP fluorescence, consistent with BG-mediated hydrolysis of cellobiose into glucose, which supported cellular growth and activated the inverted glucose sensor. Time-course measurements over 24 hours showed a graded fluorescence response proportional to cellobiose concentration. Using a fluorescence–glucose calibration curve (R² = 0.91), glucose production was estimated. 100 mM cellobiose produced approximately 12 mM glucose after 18 hours (Figure S10). These results demonstrate a closed-loop system in which a single strain converts cellobiose to glucose and reports intracellular metabolite availability via CRISPRi regulation (Figure 5E). Overall, these results illustrate the potential for real-time monitoring of metabolic processes in microbial cultures.

### Modular two-plasmid architecture enhances glucose production and sensing

To improve modularity and sensing performance, while reducing the metabolic burden of a single plasmid, we designed a two-plasmid system that separates enzymatic conversion from sensing functions (Figure 6A). A high-copy plasmid (pMB1) encodes secretion of β-glucosidase (BG), which is induced by cellobiose under the control of the mtrC promoter, CAP-regulated gRNA expression, while a low-copy plasmid (p15A) carries the CRISPRi-based glucose biosensor with constitutive dCas9, and an sfGFP reporter under the control of the PJ23108.

**Figure 6.**
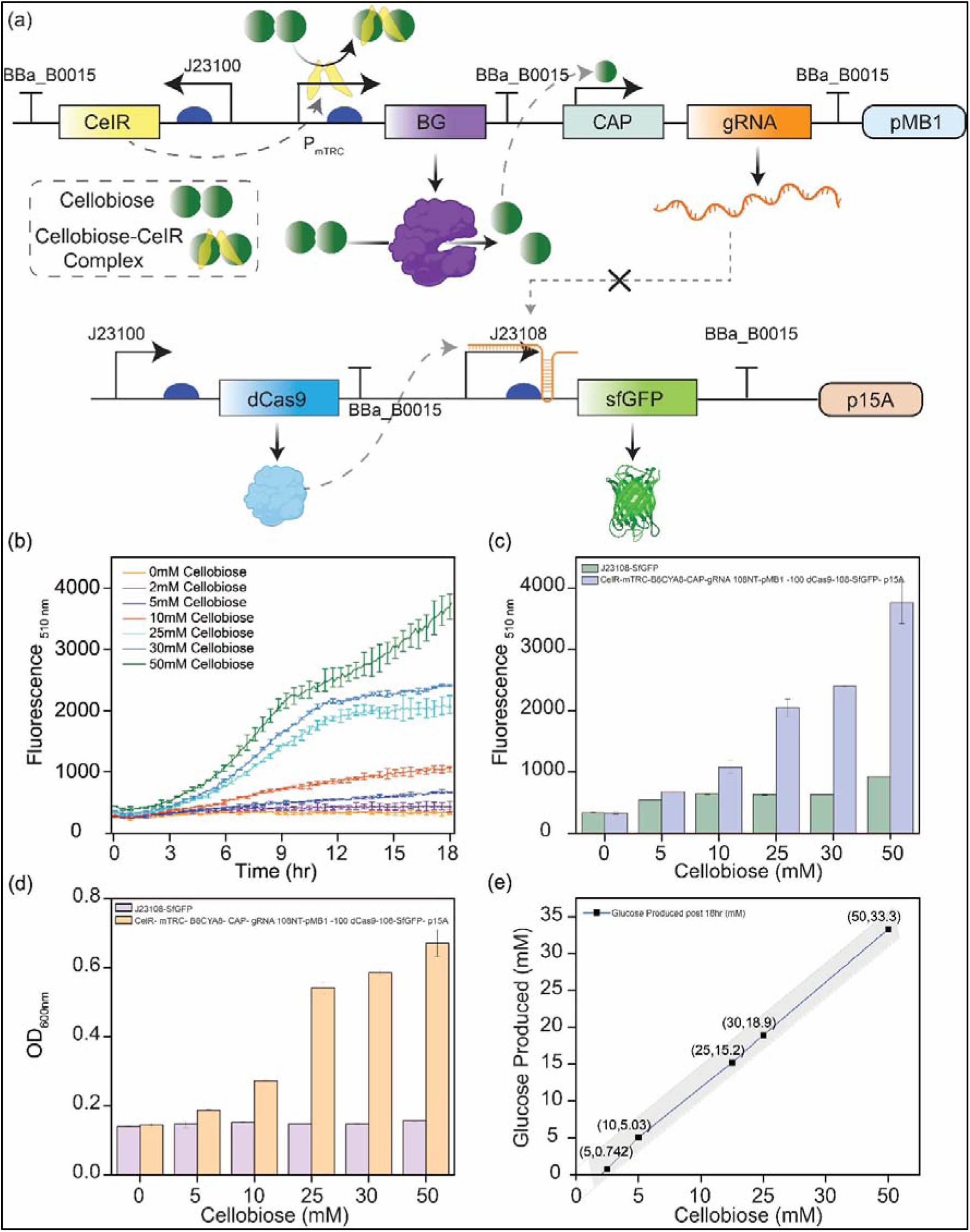
Modular two-plasmid system for glucose production and detection using high-copy BG and low-copy CRISPRi biosensor plasmids. (A) Genetic architecture of the system: BG is secreted and gRNA is expressed from a high-copy plasmid (pMB1), while dCas9 and sfGFP are expressed from a low-copy plasmid (p15A). (B) Time-course fluorescence shows dose-dependent increase in sfGFP signal with increasing cellobiose concentrations over 18 h. (C) Endpoint fluorescence and (D) OD□□□ measurements reveal glucose-dependent growth and sensing only in the engineered strain, not in the control. (E) Glucose concentration estimation via fluorescence calibration curve shows high linearity (R² = 0.9859), confirming efficient glucose production from cellobiose.

In M9 medium supplemented with 0–50 mM cellobiose, the control strain containing only the sensor plasmid showed no growth nor fluorescence induction (Figures 6C, 6D), confirming that *E. coli* cannot use cellobiose natively. In contrast, the co-transformed strain showed robust, cellobiose-dependent increase in optical density (OD_600_) and sfGFP fluorescence. Fluorescence increased linearly over time and across substrate concentrations (Figure 6B), with endpoint measurements after 18 hours exhibiting concentration-dependent differences in fluorescence intensity (Figure 6C). Glucose quantification using a fluorescence-based calibration curve (R² = 0.9859) (Figure S11) revealed that up to ∼ 33.3 mM glucose was produced from 50 mM cellobiose within the same time period (Figure 6E).

Thus, compared to the single-plasmid construct, the two-plasmid system produced more glucose, greater fluorescence intensity, and a broader dynamic range. This improvement may be attributed to modularization, which allows for high-level BG expression from a high-copy plasmid while maintaining sensor sensitivity on a low-copy backbone, thereby reducing transcriptional interference and metabolic burden. While the single-plasmid system demonstrated proof-of-concept glucose biosensing from cellobiose, the two-plasmid design offers superior scalability, tunability, and robustness, making it more suitable for quantitative biosensing and metabolic monitoring applications.

## Conclusions

In this study, we developed a synthetic, modular, and programmable whole-cell biosensor capable of detecting glucose with high specificity and dynamic responsiveness through CRISPR interference (CRISPRi). By inverting the native repression logic of a CAP-sensitive promoter with a dCas9–gRNA regulatory module, we transformed glucose-mediated repression into a glucose-inducible signal. This innovation enables tunable, real-time fluorescent output in *E. coli*. Additionally, we demonstrated the versatility of this system by coupling it with a secretory β-glucosidase (BG) enzyme to create a self-contained metabolic-sensing platform. The engineered strain effectively hydrolyzes cellobiose into glucose, supports growth in minimal media, and simultaneously reports intracellular glucose accumulation via the CRISPRi-regulated reporter. Comparative analysis of single- and dual-plasmid architectures demonstrated that functional decoupling of enzyme production from sensing circuits significantly enhances signal resolution, dynamic range, and glucose output, with the two-plasmid system yielding up to ∼33 mM glucose from 50 mM cellobiose. This work highlights the potential of CRISPRi logic reprogramming to design custom biosensors that address the limitations of native regulatory architectures. The modularity and portability of our system make it adaptable to a wide range of metabolites and regulatory inputs, offering a scalable framework for dynamic control and real-time monitoring in fields such as synthetic biology, metabolic engineering, and bioprocess optimization. Future efforts could expand this approach to include multiplexed sensing, closed-loop feedback regulation, or point-of-care diagnostics using field-deployable microbial systems.

## ASSOCIATED CONTENT

Supporting Information.

## Author Contributions

S.D. supervised the project. M.G., A.D., and S.D. conceived and designed the project. M.G. and A.D. designed all the experiments, cloned and assembled constructs, performed assays, western blots, collected and analyzed the characterization data. S.P. helped in initial experiments and figure preparation. The manuscript was written through the contributions of M.G., A.D., and S.D. All authors have given approval to the final version of the manuscript.

## Notes

The authors declare no competing financial interests.

## ACKNOWLEDGMENT

MG acknowledges DST-INSPIRE for a Senior Research Fellowship. SD acknowledges SERB Core Research Grant (CRG/2023/002111) and DBT, Government of India (grant number BT/PR47801/BCE/8/1812/2023). AD acknowledges ANRF for Junior research fellowship. The authors thank the infrastructural facilities supported by IISER Kolkata and DST-FIST (SR/FST/LS-II/2017/93(c)).

