## Supplementary Files for "Engineering a Glucose-Inducible Whole-Cell Biosensor via CRISPRi-Based Promoter Reprogramming"

**Table of Contents**

**Supplementary Figures**

Supplementary Figure S1: Fluorescence Kinetics Assay of the library of gRNAs……………3

Supplementary Figure S2: Characterization of an IPTG-Inducible Reporter Cassette with gRNA Targeting the Inducible Promoter………………………………………………………4

Supplementary Figure S3: Characterization of Cellobiose Inducible Regulator CelR………..5

Supplementary Figure S4: Quantification of BG via fluorescence co-expression studies…….6

Supplementary Figure S5: Western blot for secretion of BG with differential cellobiose induction……………………………………………………………………………………….7

Supplementary Figure S6: Characterization of gRNA 108 T 5’-3’ with Glucose responsive unit……………………………………………………………………………………………..8

Supplementary Figure S7: Characterization of gRNA 108 T 3’-5’ with Glucose responsive unit……………………………………………………………………………………………..9

Supplementary Figure S8: Biosensor Parameters for Glucose Sensing genetic circuit……...10

Supplementary Figure S9: Correlation between Fluorescence Increase rate vs Glucose consumption rate.…………………………………………………………………………….11

Supplementary Figure S10: Response of Cellobiose responsive One plasmid system in Glucose……………………………………………………………………………………….12

Supplementary Figure S11: Response of Cellobiose responsive Two plasmid system in Glucose……………………………………………………………………………………….13

**Supplementary Tables**

Supplementary Table S1: Strains and Plasmids……………………………………………...14

Supplementary Table S2: Primers……………………………………………………………16

Supplementary Table S3: Sequence of gRNAs…………………………………………………...18

Supplementary Table S4: Glucose Consumption Rate vs Fluorescence Increase Rate…….....19

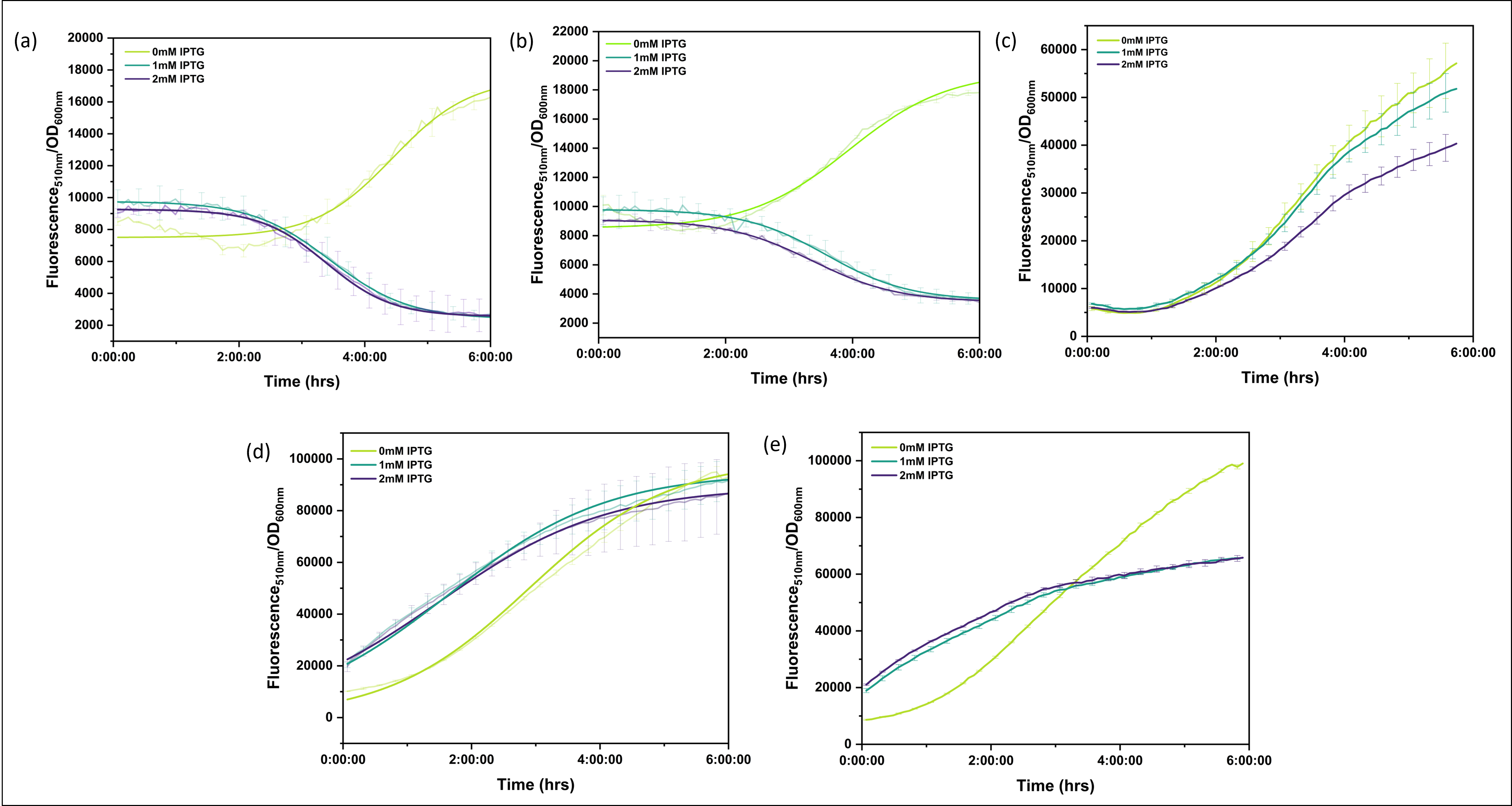

**Figure S1.** Fluorescence_510nm_/OD_600nm_ kinetics data over a 6 hour time period of the gene circuit cassette T7- gRNA-Ter-J23100-pETRBS-dCas9-Ter-Target Pro-pETRBS-sfGFP-Ter-pTU2a. The different kinetics data correspond to different orientations, target sites, and target promoters of the gRNA. (a) Fluorescence_510nm_ by OD_600nm_ Kinetics of gRNA which targets the Template region of PJ23108 with the dCas9 handle towards 3’ position and in upright orientation. (b) Fluorescence_510nm_ by OD_600nm_ Kinetics of gRNA which targets the Template region of PJ23108 with the dCas9 handle towards 5’ position and in upright orientation. (c) Fluorescence_510nm_ by OD_600nm_ Kinetics of gRNA which targets the non-template region of PSJM910 with the dCas9 handle towards the 3’ position and in inverted orientation. (d) and (e) Fluorescence_510nm_ by OD_600nm_ Kinetics of gRNA which targets the non-template region of coding sequence of sfGFP, particularly Thr65-Phe71 and Gly67-Arg73 respectively, with the dCas9 handle towards 3’ position and in inverted orientation. The cultures are induced with differential IPTG concentration, as shown in the respective graphs[1].

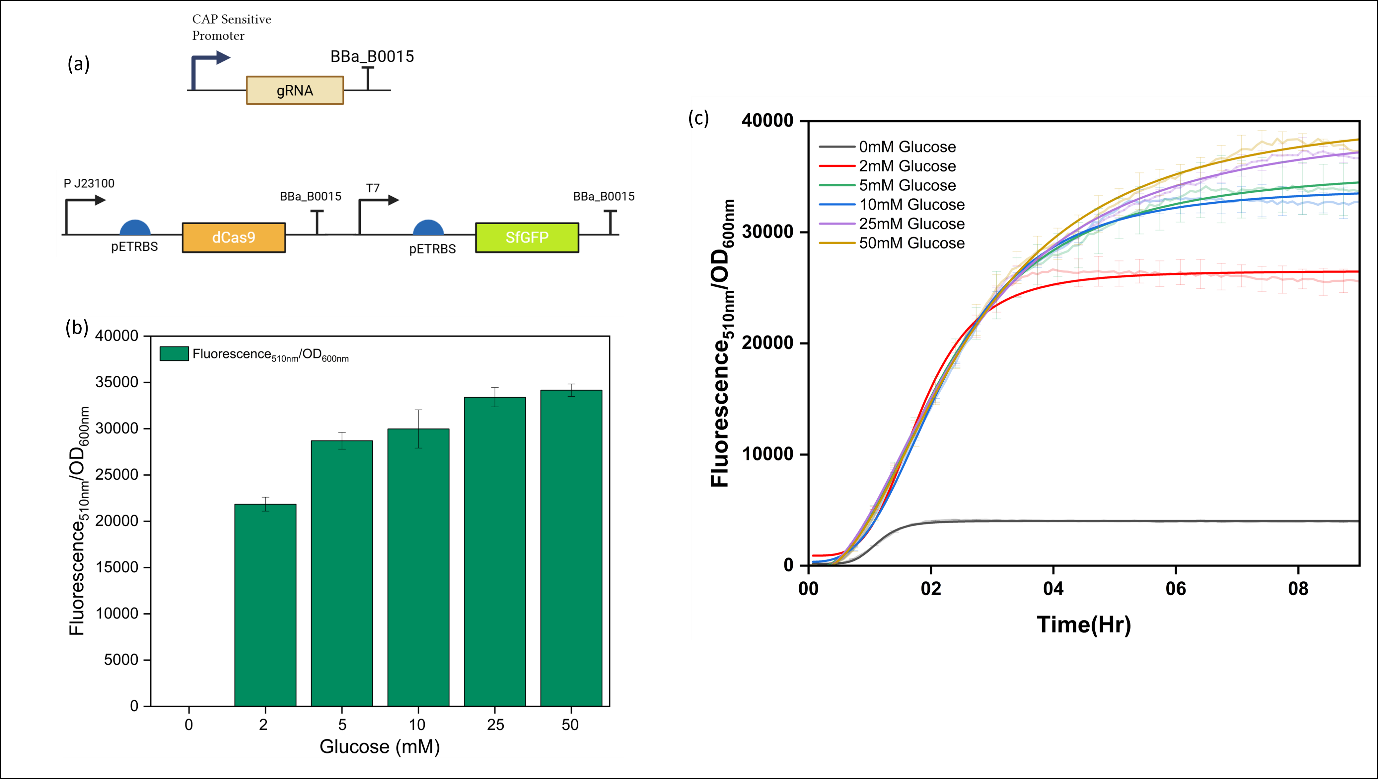

**Figure S2**. (a) Schematic of the synthetic circuit: gRNA targeting the –10 region of the IPTG inducible T7 promoter is expressed from a CAP-sensitive promoter, dCas9 is constitutively expressed, and sfGFP is under J23108 control. (b) Fluorescence response (RFU/OD_600_) across increasing glucose concentrations (0–50 mM). (c) Time-course fluorescence measurement over 8 hours at varying glucose concentrations. Reporter expression increases in a dose-dependent manner with glucose. Error bars represent standard deviation from biological triplicates.

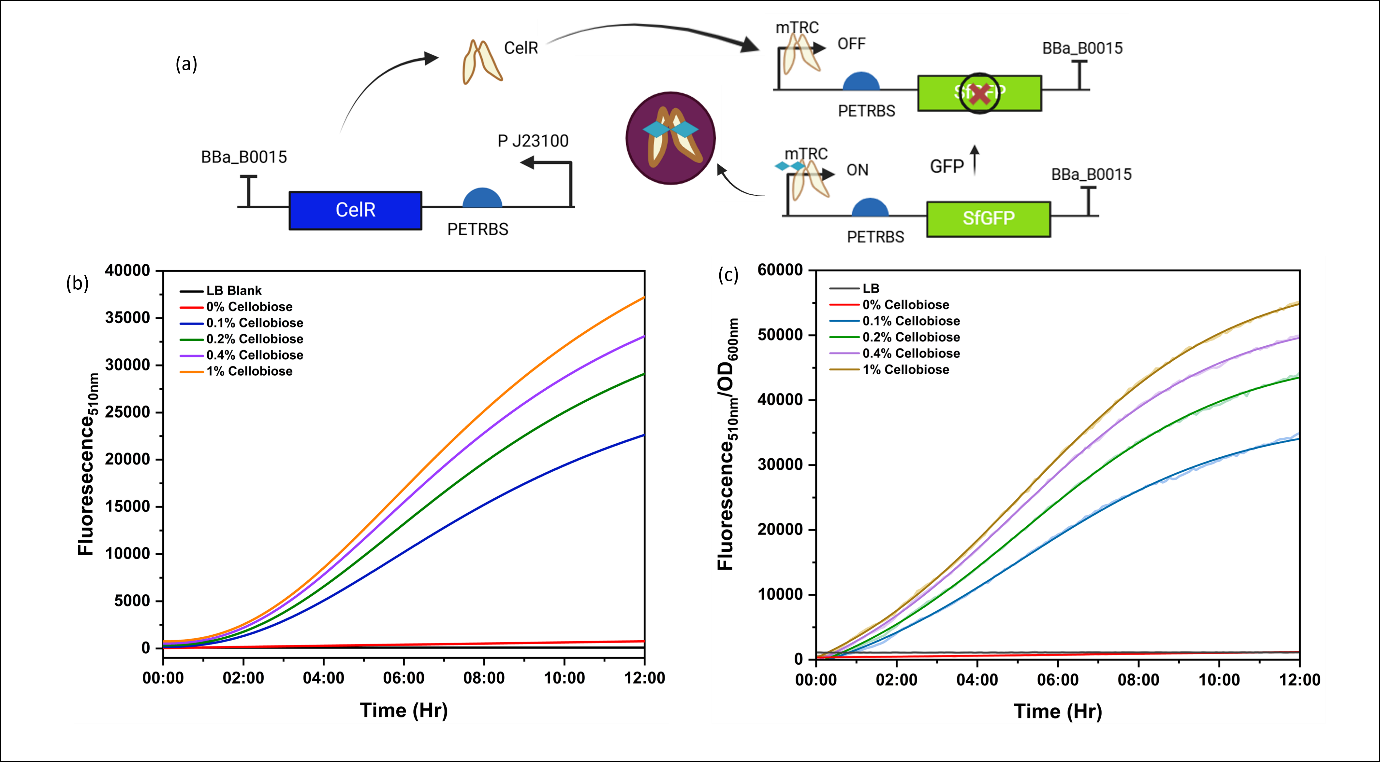

**Figure S3.** (a) Schematic of the gene circuit: Cellobiose regulator CelR under constitutive promoter, fluorescent reporter sfGFP under cellobiose induced mTRC promoter. CelR represses the expression in the absence of Cellobiose. In the presence of Cellobiose, CelR-cellobiose induces the sfGFP expression. (b) Time course Fluorescence_510nm_ measurement over 12hr time period showing a dose-dependent fluorescence response to Cellobiose. (c) Time course Fluorescence_510nm_/OD_600nm_ studies over 12hr period, indicating a strong correlation between cellobiose inducible expression of fluorescence, nullifying the dependency on growth[2].

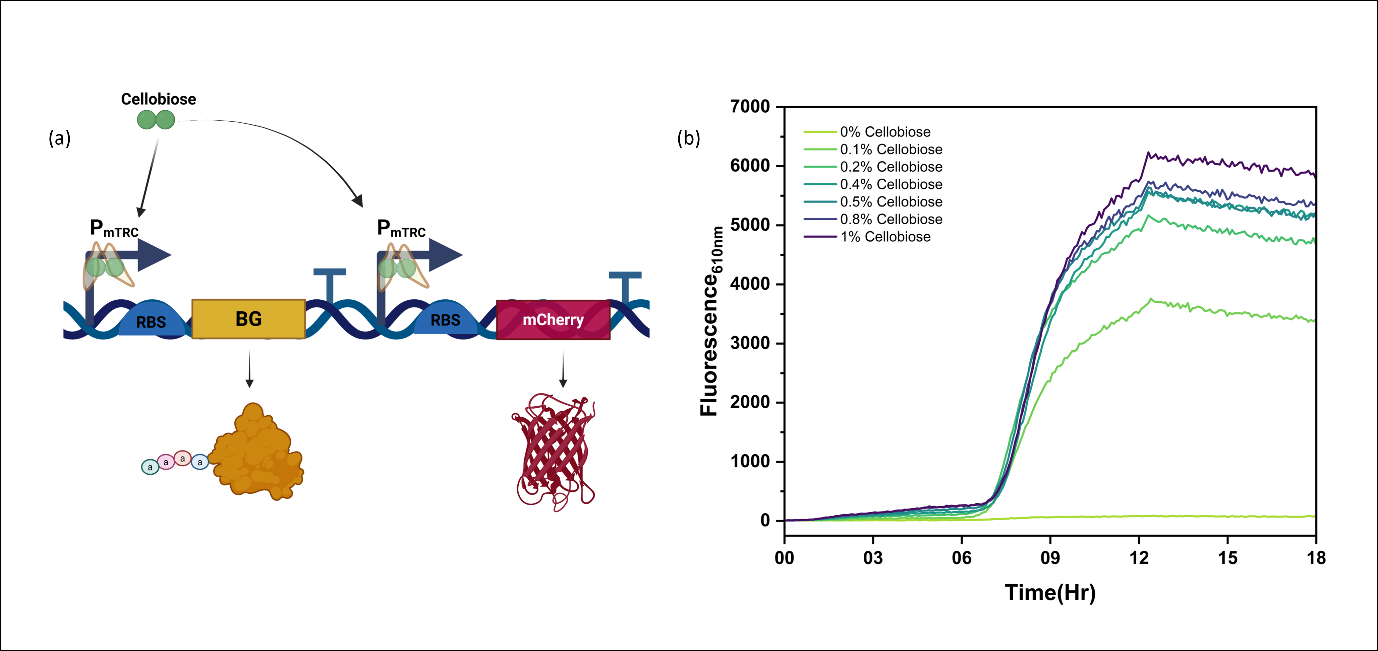

**Figure S4.** (a) Schematic of the gene circuit co-expressing β-glucosidase and mCherry under Cellobiose inducible P_mTRC_ promoter to quantify production of β-glucosidase. (b) Time course Fluorescence _610nm_ measurement over 18 hours showing a dose dependent fluorescence response to Cellobiose, indicating more β-glucosidase production under higher Cellobiose induction. mCherry production shoots up exponentially post 6 hours cellobiose induction.

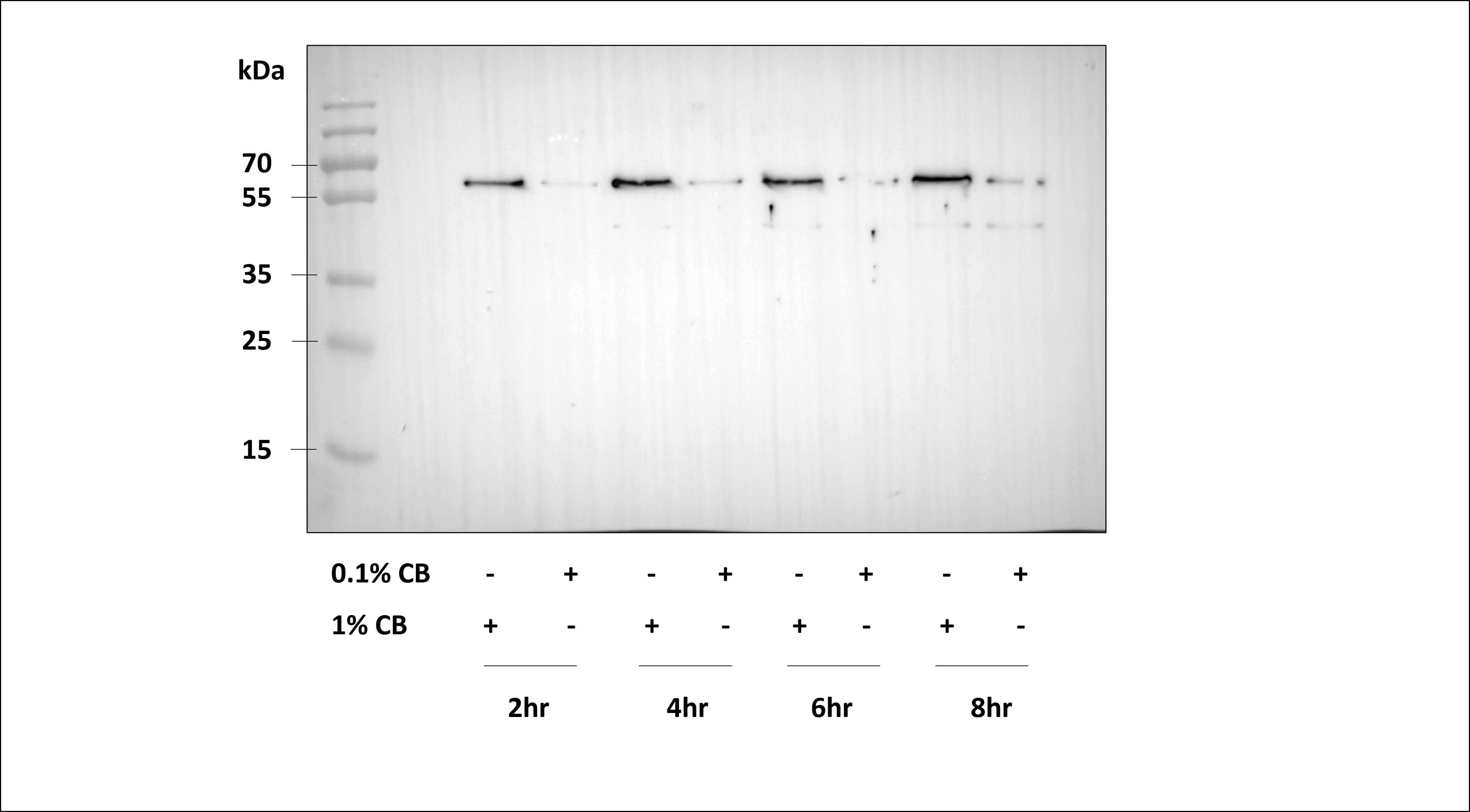

**Figure S5**. Raw Western Blot image corresponding to the differential BG secretion into the media at varying time points under varying cellobiose induction. This corresponds to Figure 4C in the main manuscript.

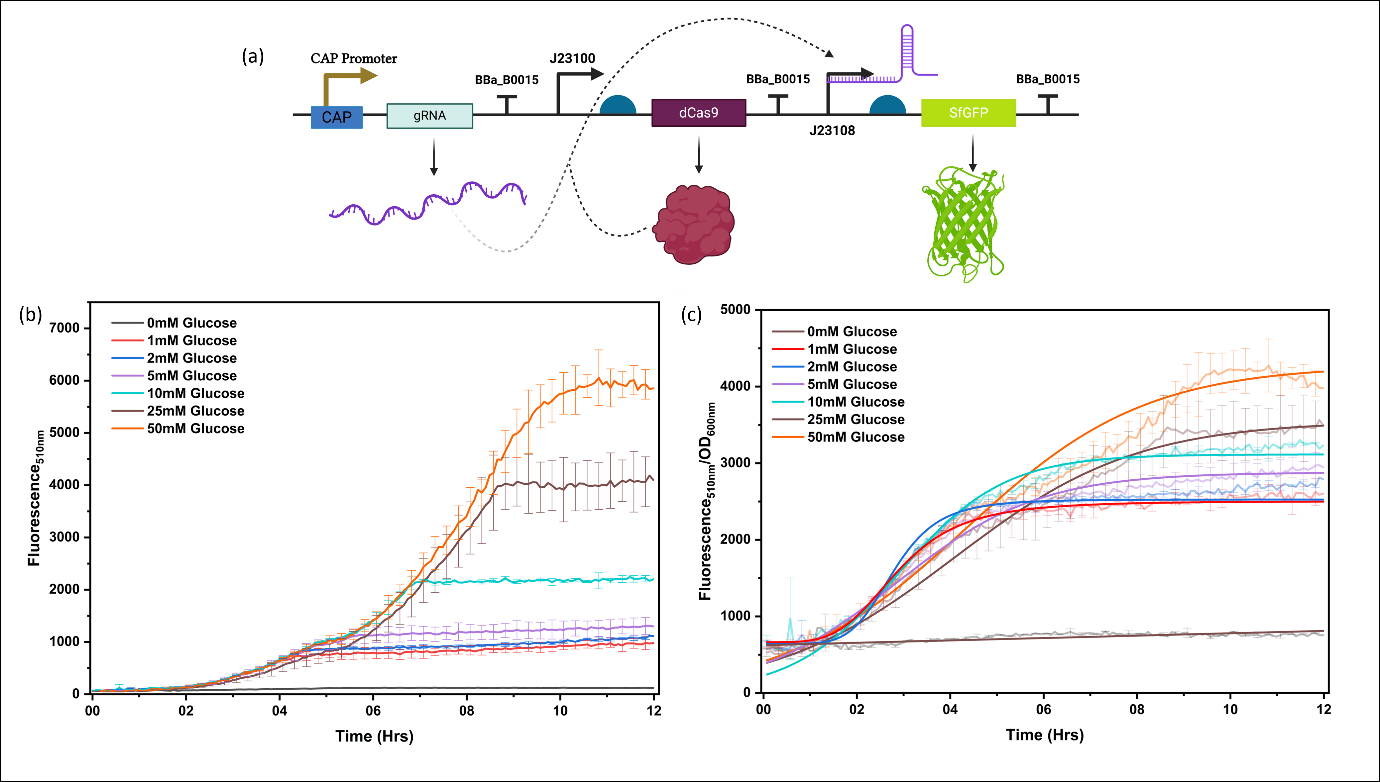

**Figure S6.** (a) Schematic of the gene circuit expressing gRNA targeting the Template region of PJ23108 with the dCas9 handle towards the 5’ position and in upright orientation under the glucose-sensitive CAP promoter, dCas9 under a constitutive promoter, and sfGFP under PJ23108. Under low-glucose conditions, less fluorescence is produced, and vice versa under high-glucose conditions. (b) Dose-dependent time course fluorescence response to varying Glucose concentrations in the growth medium. (c) Time course Fluorescence_510nm_/OD_600nm_ response of the genetic circuit over 12 hours, indicating a lower dynamic range of sensing.

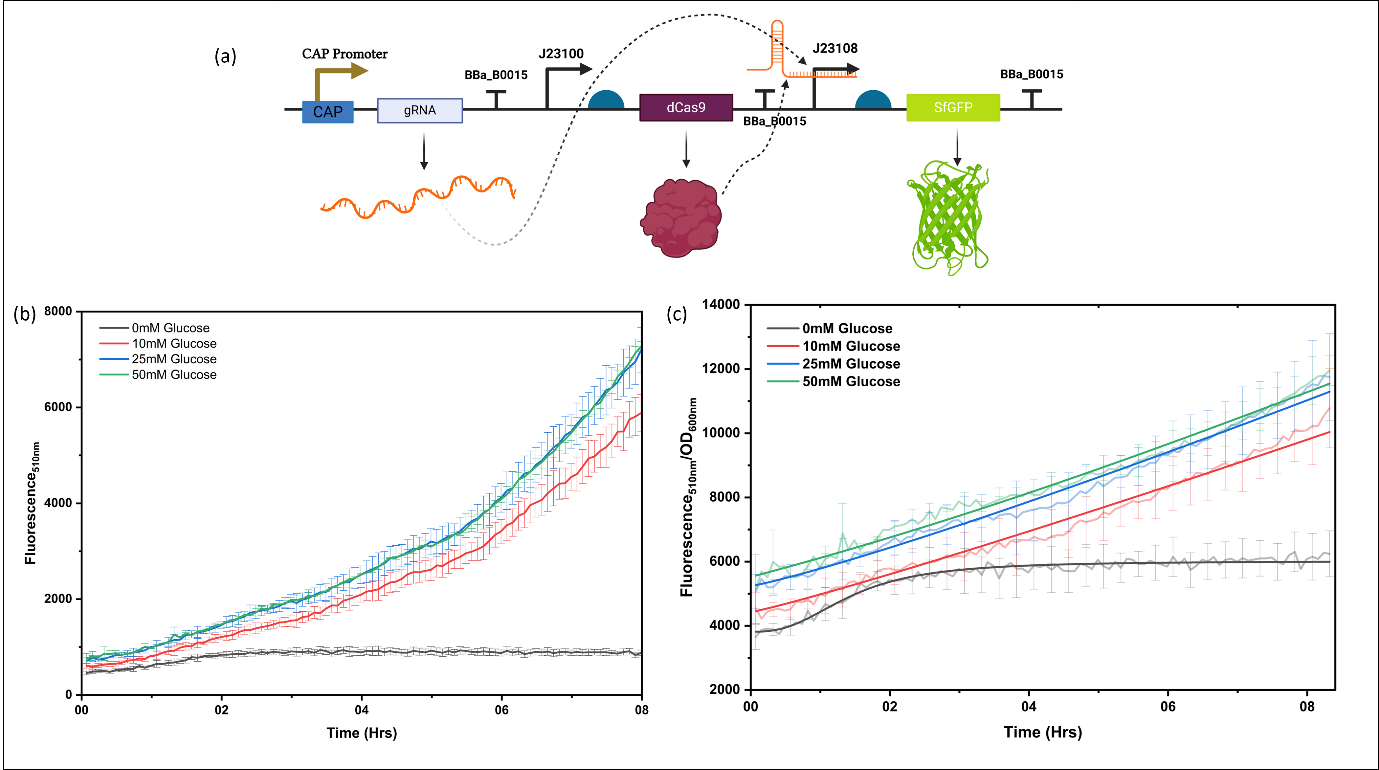

**Figure S7.** (a) Schematic of the gene circuit expressing gRNA targeting the Template region of PJ23108 with the dCas9 handle towards 3’ position and in upright orientation under Glucose sensitive CAP promoter, dCas9 under constitutive promoter and sfGFP under PJ23108. (b) Dose dependent time course fluorescence response to varying Glucose concentrations in the growth medium. (c) Time course Fluorescence_510nm_/OD_600nm_ response of the genetic circuit over 8 hours, indicating a lesser dynamic range of sensing.

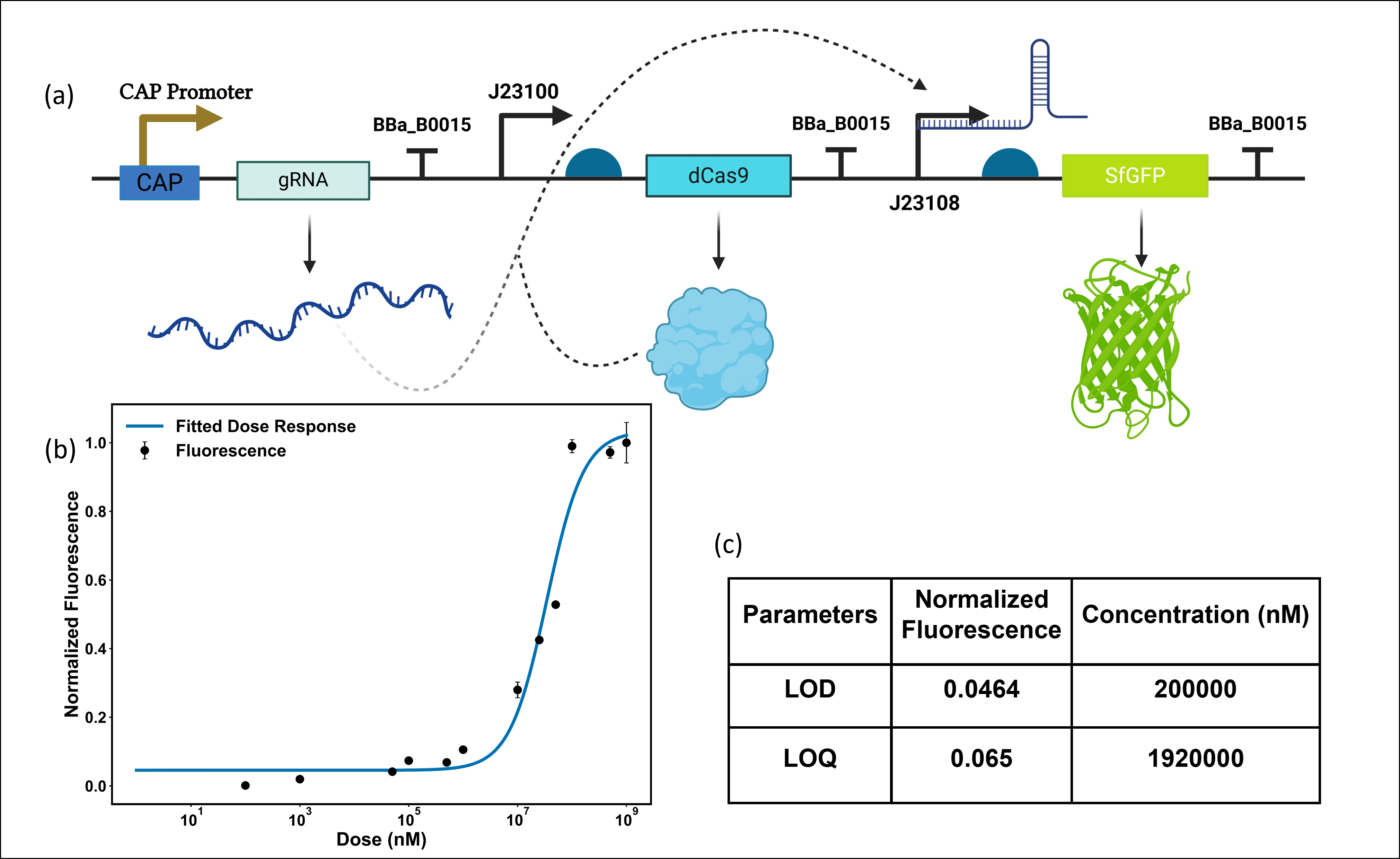

**Figure S8.** (a) Schematic of the gene circuit expressing gRNA targeting the Non-template region of PJ23108 with the dCas9 handle towards 3’ position and in inverted orientation (108 NT 5’-3’) under the glucose-sensitive CAP promoter, dCas9 under a constitutive promoter and sfGFP under PJ23108. (b) Dose Response fluorescence response to varying Glucose concentrations in the growth medium, from 100 nM-1 M concentration range. (c) The biosensor characterization parameters indicate the minimum detection range. LOD: Limit of Detection; LOQ: Limit of Quantification.

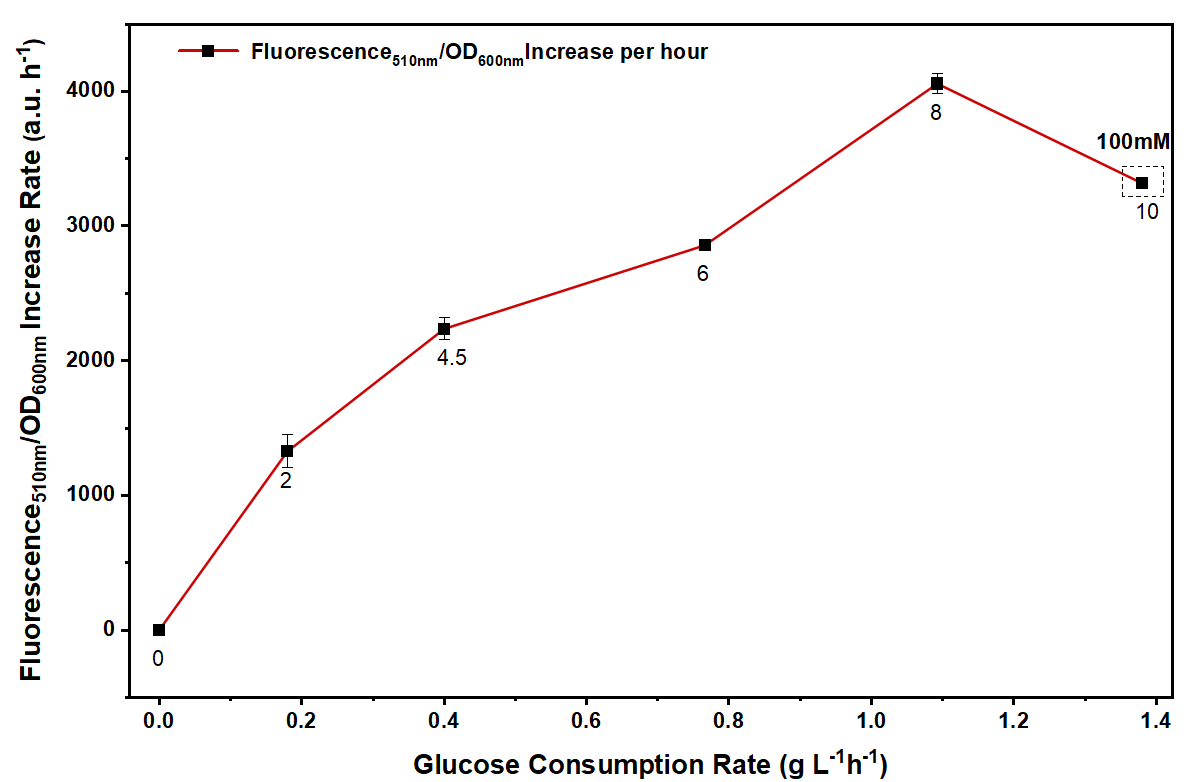

**Figure S9.** Correlation between Fluorescence_510nm_/OD_600nm_ Increase rate (per h) vs Glucose Consumption Rate (per h); Linear correlation till 50mM Glucose and a plateau with decreasing trend post 50mM till 100mM Glucose (last data point), indicating minimal efficiency in sensing post 50mM glucose concentrations (Table S4). Number annotations beside each data point indicate the time point of media collection. All assays are performed in triplicate, and the error bars indicate standard deviation.

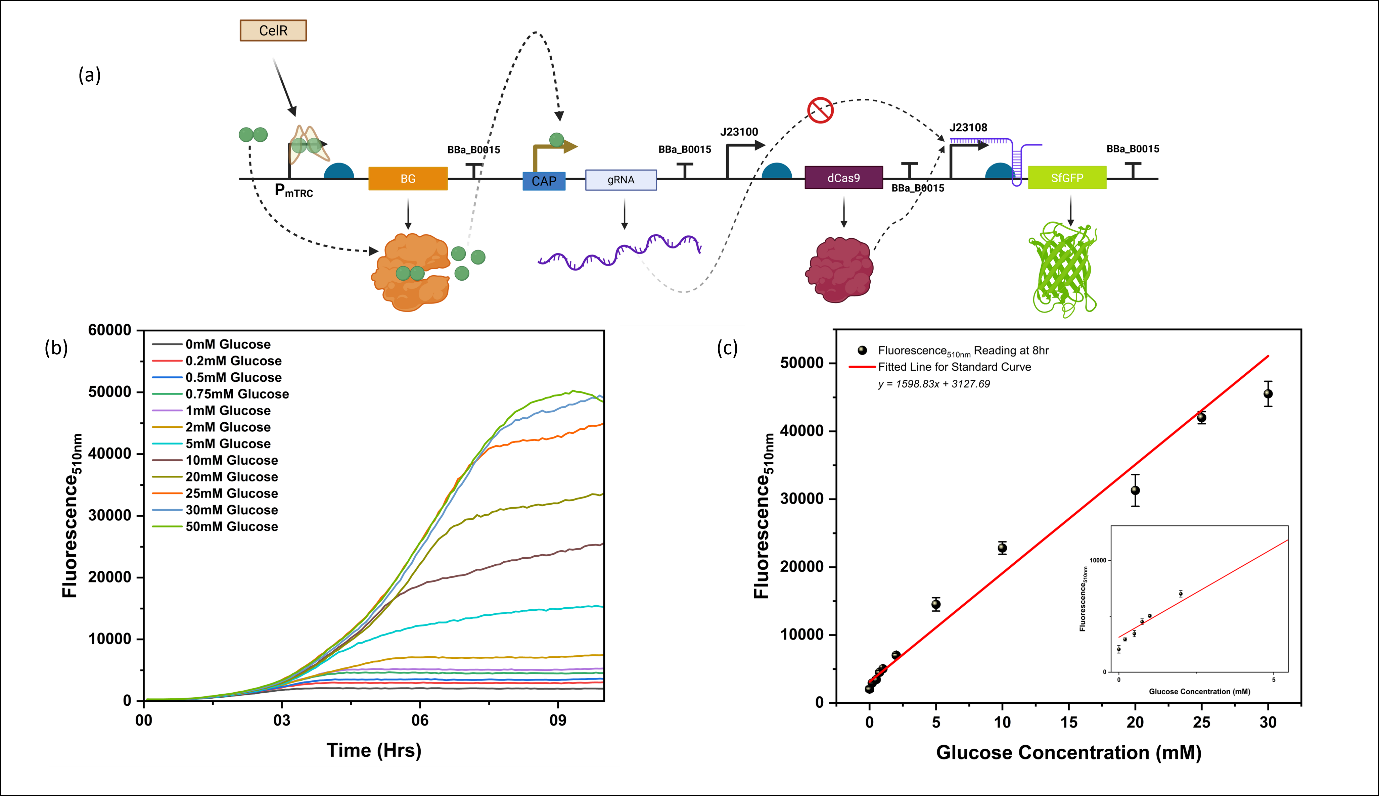

**Figure S10.** (a) Schematic of the gene circuit expressing gRNA targeting the non-template region of PJ23108 with the dCas9 handle towards 3’ position and in inverted orientation (108 NT 5’-3’) under Glucose sensitive CAP promoter, dCas9 and CelR under a constitutive promoter, and sfGFP under PJ23108, BG under P_mTRC_. (b) Dose-dependent time course fluorescence response of the cellobiose-sensitive genetic circuit to varying Glucose concentrations in the growth medium. (c) Fluorescence_510nm_ vs Glucose (mM) Standard curve indicating a linear correlation between doses of glucose and fluorescence. This standard curve is used for the quantification of cellobiose conversion to glucose in a single plasmid cassette in (a).

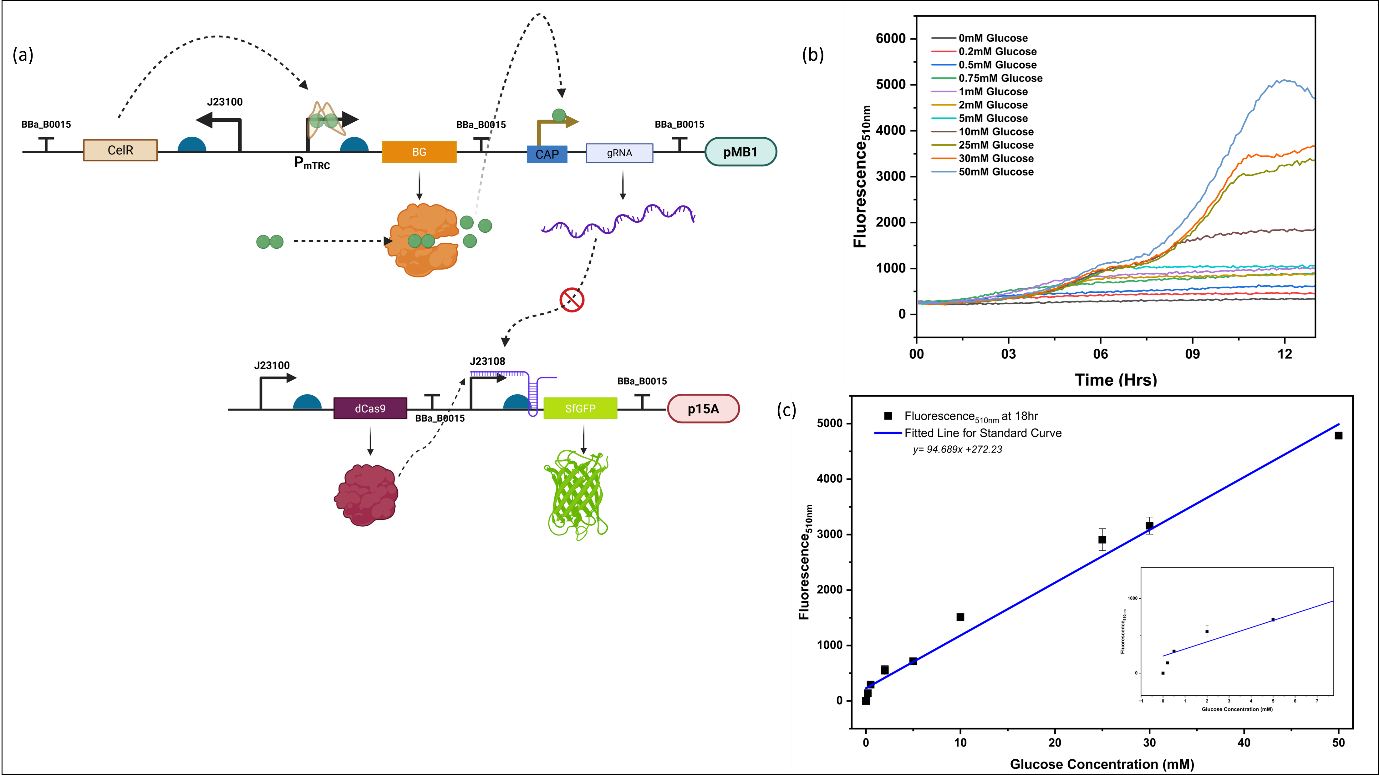

**Figure S11.** (a) Schematic of the gene circuit expressing gRNA targeting the non-template region of PJ23108 with the dCas9 handle towards 3’ position and in inverted orientation (108 NT 5’-3’) under Glucose sensitive CAP promoter, CelR under a constitutive promoter, BG under P_mTRC_ in pMB1, Chl plasmid backbone, and dCas9 under PJ23100, sfGFP under PJ23108 in p15A, Amp backbone for co-transformation. (b) Dose-dependent time course fluorescence response of the cellobiose-sensitive genetic circuit to varying Glucose concentrations in the growth medium. (c) Fluorescence_510nm_ vs Glucose (mM) Standard curve indicating a linear correlation between doses of glucose and fluorescence. This standard curve is used for the quantification of cellobiose conversion to glucose in the dual plasmid cassette in (a).

**Table S.** Strains and plasmids used in this study

| **Strains or Plasmids** | **Characteristics** | **References** |
| --- | --- | --- |
| **Strains** |  |  |
| DH5α | F−∆lacU169(Ø80d lacZ∆M15) supE44 hsdR17 recA1 gyrA96 endA1 thi-1 relA1 |  |
| BL21(DE3) |  |  |
| **Plasmids** |  |  |
| pBP- J23100 | Constitutive Anderson Promoter | **[3]** |
| pBP-PET RBS | PET RBS | **[3]** |
| pBP-J23108 | Constitutive Anderson Promoter | **[3]** |
| pBP-T7 | T7 promoter | **[3]** |
| pBP-Tag_linker | PET RBS with tag linker | **[3]** |
| pBP-lacz | Vector to carry Promoter in place of lacZ | **[3]** |
| pBP-ORF | Empty vector to clone ORF | **[3]** |
| pBP-Bba_B0015 | Synthetic double terminator | **[3]** |
| pBP-BG | β-glucosidase at level 0 | **[3]** |
| pBP-lacZ-ansb | ansB tag at level zero | **[3]** |
| pBP-CAP | CAP Sensitive Promoter at level 0 | This work |
| pBP-mTRC | Cellobiose Regulated Promoter with CelR | This work |
| pBP-sfGFP | Super folder Green Fluorescent Protein (sfGFP) at level 0 | This work |
| pBP-CelR | Transcriptional Regulator of Cellobiose Induction at level 0 | This work |
| pBP-gRNA -10/108 NT | gRNA targeting J23108 -10 Template region at level 0 | This work |
| pBP-dCas9 | dead Cas9 protein at level 0 | This work |
| pTU1A- CAP-sfGFP | sfGFP under CAP Sensitive Promoter, AmpR | This work |
| pTU1A- J23100-CelR | CelR under J23100 Promtoter at level 1, AmpR | This work |
| pTU1B-mTRC-sfGFP | sfGFP under Cellobiose Sensitive mTRC Promoter, AmpR | This work |
| pTU1B-mTRC-ansb B8CYA8 | β-glucosidase with ansb tag at level 1 under mTRC Promoter, AmpR | This work |
| pTU1D-CAP-gRNA -10/108 NT | gRNA targeting J23108 -10 Template region under CAP Promoter at level 1, AmpR | This work |
| pTU1A-J23100-dCas9 | dCas9 protein under J23100 promoter at level 1, AmpR | This work |
| pTU1A-J23108-sfGFP | sfGFP under J23108 at level 1, AmpR | This work |
| pTU1D-CAP-gRNA -10/910 NT | gRNA targeting SJM910 -10 Template region under CAP Promoter at level 1, AmpR | This work |
| pTU1D-CAP-gRNA -10/T7 NT | gRNA targeting T7 -10 Template region under CAP Promoter at level 1, AmpR | This work |
| pTU2a-CelR-mTRC-sfGFP | CelR and sfGFP (under mTRC) assembled in level 2, ChlR | This work |
| pTU2a-dCas9-J23108-sfGFP | dCas9 and J23108-sfGFP assembled in level 2, ChlR | This work |
| pTU2a-T7-gRNA -10/108 NT- J23100-dCas9-J23108-sfGFP | gRNA targeting J23108 -10 Non Template region under T7, dCas9 under J23100 and sfGFP under J23108 at level 2, ChlR | This work |
| pTU2a-T7-gRNA -10/910 NT- J23100-dCas9-J23108-sfGFP | gRNA targeting SJM910 -10 Non Template region under T7, dCas9 under J23100 and sfGFP under SJM910 at level 2, ChlR | This work |
| pTU2a-T7-gRNA -10/108 T 5'-3'- J23100-dCas9-J23108-sfGFP | gRNA targeting J23108 -10 Template region under T7, dCas9 (with handle towards 3' end) under J23100 and sfGFP under J23108 at level 2, ChlR | This work |
| pTU2a-T7-gRNA -10/108 T 3'-5'- J23100-dCas9-J23108-sfGFP | gRNA targeting J23108 -10 Template region under T7, dCas9 (with handle towards 5' end) under J23100 and sfGFP under J23108 at level 2, ChlR | This work |
| pTU2a-T7-gRNA -Gly66- J23100-dCas9-J23108-sfGFP | gRNA targeting sfGFP Gly67-Arg73 Template region under T7, dCas9 (with handle towards 5' end) under J23100 and sfGFP under J23108 at level 2, ChlR | This work |
| pTU2a-T7-gRNA -Thr65- J23100-dCas9-J23108-sfGFP | gRNA targeting sfGFP Thr65-Phe71 Template region under T7, dCas9 (with handle towards 5' end) under J23100 and sfGFP under J23108 at level 2, ChlR | This work |
| pTU2a-CAP-gRNA -10/108 NT- J23100-dCas9-J23108-sfGFP | gRNA targeting J23108 -10 Template region under CAP, dCas9 under J23100 and sfGFP under J23108 at level 2, ChlR | This work |
| pTU2a-CAP-gRNA -10/T7 NT- J23100-dCas9-T7-sfGFP | gRNA targetingT7 -10 Template region under CAP, dCas9 under J23100 and sfGFP under T7 at level 2, ChlR | This work |
| pTU2a-CAP-gRNA -10/108 T 3'-5'- J23100-dCas9-J23108-sfGFP | gRNA targeting J23108 -10 Template region under CAP, dCas9 (with handle towards 5' end) under J23100 and sfGFP under J23108 at level 2, ChlR | This work |
| pTU2a-CAP-gRNA -10/108 T 5'-3'- J23100-dCas9-J23108-sfGFP | gRNA targeting J23108 -10 Template region under CAP, dCas9 (with handle towards 3' end) under J23100 and sfGFP under J23108 at level 2 | This work |
| pTU2A(ColE1)-CelR-mTRC-B8CYA8 | CelR and ansb-B8CYA8 (under mTRC) assembled in level 2 vector 2A ColE1 ori, ChlR | This work |
| pTU2A(ColE1)-CelR-mTRC-B8CYA8-mTRC-mCherryssrA | CelR and ansb-B8CYA8 (under mTRC) and mCherry (under mTRC) assembled in level 2 vector 2A ColE1 ori, ChlR | This work |
| pTU2a-CelR-mTRC-B8CYA8-CAP-gRNA -10/108 NT-J23100-dCas9-J23108-sfGFP | CelR, ansb-B8CYA8, CAP gRNA -10/108 NT, dCas9 under J23100 and sfGFP under J23108 in level 2 pTU2a construct, ChlR | This work |
| pTU2a-CelR-mTRC-B8CYA8-CAP-gRNA -10/108 NT | CelR, ansb-B8CYA8, CAP gRNA -10/108 NT in level 2 pTU2a construct, ChlR | This work |
| pTU2A (p15A)-J23100-dCas9-J23108-sfGFP | dCas9 under J23100 and sfGFP under J23108 in level 2 pTU2A (p15A ori) construct, AmpR | This work |

**Table S2.** Primers used in this study

| **Primer** | **DNA Sequences** |
| --- | --- |
| CAP_FP | GGAATTCCATATGGGTCTCACTATTAATGTGAGTTAGCTCACTCATTAGGC |
| CAP_RBS_RP | CTGCAGCATGCGGTCTCTTATGTTTCTCCTCTTTCTCTAGTATGTGTGAAATTG |
| CAP RP | GGCATGCGGTCTCTTATGCTCTAGTATGTGTGAAATTGTTATCCGC |
| CelR_FP | GGTCTCACATATGGAGCGTCG |
| CelR_RP | GGTCTCTTCGATTAGTGGTGGTGG |
| trc_Gibson_P1_RP | GCGCTCCCACCACACATTATACGAGCCGGATGATTAATTGTCAAATAGTGAGACCCATATGCGTCTCTAGATTCTAGAAGCGGC |
| trc_Gibson__P2_FP | CGTATAATGTGTGGTGGGAGCGCTCCCATCACACAGGAAACAGGTACAGAGACCGCATGCCGTCTCATTAGCTGC |
| sfGFP FP | GTATGGTCTCACATATGAGCAAAGGAGAAGAACTTTTCACTG |
| sfGFP RP | ATTAGGTCTCTTCGATTATTTGTAGAGCTCATCCATGCCATG |
| ansb FP | GTATGGTCTCATAAATGGAGTTTTTCAAAAAGAC |
| ansb RP | CACATGGTCTCTTATGTGCCAATGCTGCACCAC |
| B8CYA8 FP | GGTCTCACATAATGGCGAAGATTATTTTTCCAG |
| B8CYA8 RP | GGTCTCTTCGATCAATGATGATGATGATGATGATTTGC |
| dCas9 FP | GTATGGTCTCACATAATGGATAAGAAATACTCAATAGGCTTAGCTATC |
| dCas9 RP | ATTAGGTCTCTTCGATTAGTCACCTCCTAGCTGACTCAAATC |
| gRNA -10 108 NT FP | GGGCCCACATATGGGTCTCACATAGTACGCTAGCATTATACCTAGGGTTTTAGAGCTAGAAATAGCAAGTTAAAATAAGGC |
| gRNA -10 108 NT RP | CCCGGAGCATGCTCGAAGAGACCGCACCGACTCGGTGCC |
| gRNA -10 910 NT FP | GGGCCCACATATGGGTCTCACATAGTACGCTAGCACAGTACCAAGGGTTTTAGAGCTAGAAATAGCAAGTTAAAATAAGGC |
| mCherry FP | GTATGGTCTCACATAATGGTGAGCAAGGGCG |
| mCherry ssrA RP | ATTAGGTCTCTTCGATTACGCTGCAAGGGCGTAATTTTCGTCGTTCGCTGCCTTGTACAGCTCGTC |
| CAP/T7 FP | CAGAAGCTATTAATACGACTCACTATAGGGAGAGTACTTTCACACATACTAGAGCATAGTACGCTAGC |
| CAP/T7 RP | GTGAAAGTACTCTCCCTATAGTGAGTCGTATTAATAGCTTCTGAGACGTTCTAGAAGCGGCCAGTATC |
| L4440 FP | CTTCGCGTTATGCAGGCTTCCTCGCTCACTGACTCGCT |
| L4440 | AGCGAGTCAGTGAGCGAG |
| Fragment 1 FP | TCACTATATCTTAGCATCAGTGATACTGGCCGCTTCTAGAAC |
| Fragment 1 RP | CAACTGAACATATTACCGTAGATAAGCTAGTATAGATAACAACATATAAACGCAGAAAG |
| Fragment 2 FP | GTTATCTATACTAGCTTATCTACGGTAATATGTTCAGTTGGGCCGCTTCTAGAAC |
| Fragment 2 RP | ATTACATTCGATTACGTCACCAGTAGGTCAGCCATCACGTAACATATAAACGCAGAAAG |
| Fragment 3 FP | ACGTGATGGCTGACCTACTGGTGACGTAATCGAATGTAATGGCCGCTTCTAGAAC |
| Fragment 2/3 RP | CTCTGTACCCGGAGTAGATAGATATTGATGTCGAATATAGAACATATAAACGCAGAAAG |
| 2a/2A FP | CTATATTCGACATCAATATCTATCTACTCCGGGTACAGAGACC |
| 2a/2A Gibson RP | GTTCTAGAAGCGGCCAGTATCACTGATGCTAAGATATAGTGAGACCTCTAGAAG |
| gRNA -10 mod 108 FP | GAGCATACATGCGATCGTAATATGGATCCGTTTTAGAGCTAGAAATAGCAAGTTAAAATAAGGC |
| gRNA -10 mod 108 RP | CTAAAACGGATCCATATTACGATCGCATGTATGCTCTAGTATGTGTGAAAGTACTCTCC |
| gRNA 108modified FP | GAGCATAGTCCTAGGTATAATGCTAGCGTTTTAGAGCTAGAAATAGCAAGTTAAAATAAGGC |
| gRNA 108modified RP | CTAAAACCAGGATCCATATTACGATCGTATGCTCTAGTATGTGTGAAAGTACTCTCC |
| gRNA sfGFP Gly66 FP | CATACTAGAGCATACGGGAAAAGCATTGAACACCGTTTTAGAGCTAGAAATAGCAAGTTAAAATAAGGC |
| gRNA sfGFP Gly66 RP | CTAAAACGGTGTTCAATGCTTTTCCCGTATGCTCTAGTATGTGTGAAAGTACTCTCC |
| gRNA sfGFP Thr65 FP | CATACTAGAGCATATTTCGTAACTTGTGGTATCCGTTTTAGAGCTAGAAATAGCAAGTTAAAATAAGGC |
| gRNA sfGFP Thr65 RP | CTAAAACGGATACCACAAGTTACGAAATATGCTCTAGTATGTGTGAAAGTACTCTCC |
| Vec2a Part 1 FP | TCTCACTATATCTTAGCATCAGTGATAC |
| Vec2a Part 1 RP | GATATTGATGTCGAATATAGAACATATAAACGCA |
| Vec2a Part 2 FP | GGCCTTTCTGCGTTTATATGTTCTATATTCGACATCAATATCGCATCAGTGATACTGGCCGC |
| Vec2a Part 2 RP | GGTCTCTGTACCCGGAGTAGATAGATATTGATGTCGAATATAG |

**Table S3.** gRNA Library constructed in this study

| **Promoter/Protein** | **gRNA** | **Sequence** |
| --- | --- | --- |
| J23108 | 108 NT 5’-3’ | 5’-GTACGCTAGCATTATACCTAGG-3’ |
| J23108 | 108 T 3’-5’ | 5’-CATGCGATCGTAATATGGATCC-3’ |
| J23108 | 108 T 5’-3’ | 5’-GTCCTAGGTATAATGCTAGC-3’ |
| SJM911 | 911 NT 5’-3’ | 5’-GTACGCTAGCACAGTACCAAGG-3’ |
| T7 | T7 NT 5’-3’ | 5’-CATGCGATCGTAATATGGATCC-3’ |
| sfGFP | Gly67-Arg73 NT 3’-5’ | 5’-CGGGAAAAGCATTGAACACC-3’ |
| sfGFP | Thr65-Phe71 T 3’-5’ | 5’-TTTCGTAACTTGTGGTATCC-3’ |

**Table S4.** Glucose consumption rate vs Fluorescence increase rate

| **Time (Hr)** | **Glucose consumed per hr (g L^-1^h^-1^)** | **Fluorescence510nm/OD600nm per hr (a.u. h^-1^)** | **Std. Dev.** |
| --- | --- | --- | --- |
| **0** | 0 | 0 | 0 |
| **2** | 0.18 | 1326.31 | 122.43 |
| **4.5** | 0.4 | 2236.34 | 81.73 |
| **6** | 0.77 | 2857.88 | 17.46 |
| **8** | 1.09 | 4057.95 | 74.83 |
| **10** | 1.38 | 3317.8 | 42.22 |
